# Manipulating plant oxygen sensing through NCO substitution reveals trade-offs between growth and flooding tolerance

**DOI:** 10.1101/2025.07.24.665511

**Authors:** Monica Perri, Yuming He, Ximena Chirinos, Mikel Lavilla-Puerta, Rebecca Latter, Thomas Bancroft, Antonis Papachristodoulou, Emily Flashman, Francesco Licausi

**Author notes:** **Contact info Correspondence:**. **Author Contributions**: M.P. designed and performed the research, analysed the data and wrote the manuscript. Y.H., X.C., M.L.-P. performed the research and analysed the data. R.L. analysed the data. T.B. contributed new reagents and analysed the data. A.P. designed the research. E.F. designed the research and analysed the data. F.L. designed the research, analysed the data and wrote the manuscript. **Competing Interest Statement**: The authors declare no competing interest. **Classification:** Biological sciences/Plant Biology.

## Abstract

The oxygen-dependent degradation of Ethylene Response Factors VII (ERFVIIs) through the N-degron pathway is central to regulating the transcriptional responses to hypoxia in vascular plants. Plant Cysteine Oxidases (PCOs) control this step by catalysing the oxidation of an N-terminal Cys residue exposed by ERFVIIs. In the present study, we investigated the functional impact of replacing Arabidopsis PCOs with diverse N-terminal cysteine oxidases (NCOs) from across the three eukaryotic kingdoms, hypothesizing that structural and kinetic differences may influence gene regulation of ERFVII targets under hypoxia and thus impact stress tolerance. Combining structural analyses, *in vitro* biochemical characterisation and *in planta* complementation assays we observed that not all tested NCOs are functionally equivalent to the endogenous PCOs. In fact, despite the remarkable conservation of catalytic motifs, we identified key differences in enzyme architecture that appear to affect the enzyme’s capacity to regulate hypoxia responses in plants. Notably, NCO efficiency in oxidising ERFVII peptides inversely correlated with hypoxic gene expression under aerobic conditions and enhanced submergence survival, suggesting that partial ERFVII stabilization primes plants to cope with hypoxia. However, enhanced basal expression of hypoxia-responsive genes in turn correlated negatively with development and biomass accumulation, pointing to a trade-off between growth and stress resilience. Our findings demonstrate that tuning NCO activity can reshape the transcriptional and physiological hypoxia response, suggesting it is possible to enhance plant resilience under fluctuating oxygen conditions through enzyme engineering and precision breeding.

**Significance statement:** Control of low oxygen responses to improve crop flooding tolerance is a long-sought objective of molecular plant breeders. The oxygen-dependent oxidation of N-terminal cysteines in transcription factors is thought to be a key step in modulating the transcriptional response to hypoxia. In this study, we tested this hypothesis by substituting endogenous N-terminal cysteine dioxygenases with homologues from different species characterized by highly divergent sequences, structures, and kinetic properties. We show that indeed these variations effectively and predictably influence gene transcription in plants exposed to hypoxia, thereby affecting their tolerance to submergence. However, we also demonstrate that, unexpectedly, these substitutions impact plant growth and development under aerobic conditions, revealing a trade-off between flooding stress resilience and biomass accumulation or yield.

## Introduction

Low-O_2_ responses in vascular plants rely on proteolytic control of Cys-initiating (N-cys) members of the group VII of the ethylene response factors (ERFVIIs) (1). In angiosperms, additional contribution to accommodate chronic, acute and fluctuating hypoxia is provided by the Polycomb Repressive Complex (PRC) 2 subunit Vernalization 2 (VRN2) and the LITTLE ZIPPER 2 (ZPR2) (2–4).

These N-cys proteins undergo O_2_-dependent destabilisation via post-translational regulation through the N-degron pathway (5–7). This involves Plant Cysteine Oxidases (PCOs), non-heme iron (Fe(II))-dependent dioxygenases, which act as main O_2_ sensors by catalysing oxidation of the N-terminal cysteine (8, 9). The resulting sulfinylated cysteine is recognised for arginylation by arginyltransferase (ATE), which leads to ubiquitination by the E3 Ubiquitin-ligase PROTEOLYSIS 6 (PRT6) and BIG, thereby targeting the protein to the proteasome for degradation (10, 11). Notably, N-cys degron-mediated proteolysis also requires nitric oxide (NO) (12). Oxygen depletion under hypoxic conditions prevents PCO activity, stabilising N-cys substrates, which then accumulate in the nucleus to trigger the activation of hypoxic response genes, including those coding for anaerobic metabolism (**Figure 1a**) (13, 14).

**Figure 1.**
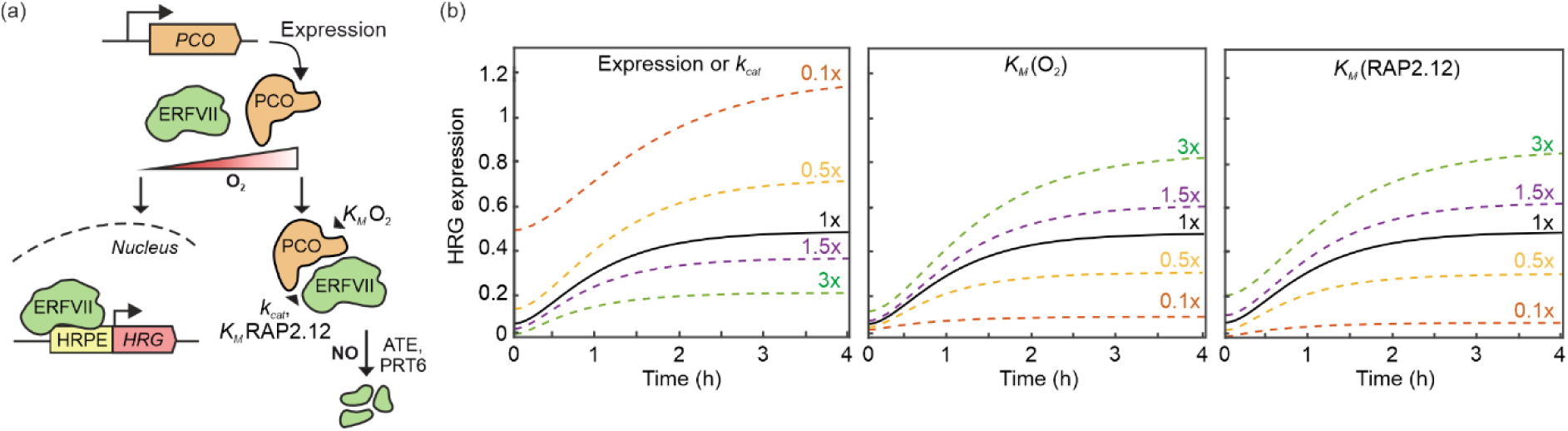
Hypoxic gene induction as a function of PCO features. (**a**) Oxygen-dependent regulation of ERFVIIs in flowering plants. ERFVII substrates are characterized by an N-terminal cysteine which, in presence of oxygen (O_2_) is targeted for oxidation by Plant Cysteine Oxidases (PCOs). The resulting Cys-sulfinic or Cys-sulfonic acid triggers proteasomal degradation of ERFVIIs. The activity of PCOs is influenced by their expression levels as well as their kinetic parameters (*K_M_* and *k_cat_*) for substrates such as RAP2.12 and O₂. In the absence of O_2_, PCOs are unable to oxidize the N-terminal cysteine, leading to stabilization of the ERFVIIs, which in turn can enter the nuclei and activate *HRG* by binding the hypoxia-responsive promoter element (HRPE). (**b**) Prediction dynamics of a generic *HRG* at 1% O_2_ (v/v) depending on PCO4 expression levels or catalytic activity (*k_cat_*), O₂ affinity (*K_M_* (O_2_)) and RAP2.12 affinity (*K_M_* (RAP2.12)) over time. Dotted lines of different colours indicate fold variation of the parameter level (black line) described in Shukla, *et al.*, 2024 (40).

The identification of PCOs in vascular plants led to the discovery that the human homolog, 2-aminoethanethiol dioxygenase (ADO), also acts as an enzymatic O_2_ sensor, alongside the Prolyl Hydroxylases (PHDs) and histone demethylases (15–17). ADO catalyses the oxidation of N-terminal cysteine residue of N-cys proteins involved in cell signalling such as the regulator of G-protein signalling 4 and 5 (RGS4/5), interleukin-32 (IL-32) and the member 10 of the acyl-CoA dehydrogenase family (ACAD10), thus modulating angiogenesis, inflammation and autophagy (17–19). Sequence alignments revealed conserved catalytic site features between PCO and ADO, despite the absence of shared targets (20, 21). Moreover, experiments *in planta* demonstrated the ability of *Homo sapiens* ADO (HsADO) to complement the homozygous quadruple *pco1/pco2/pco4/pco5* (*4pco*) mutant (17). A recent study using an *in vitro* reporter assay also confirmed that HsADO can turn over ERFVII peptides, alongside those corresponding to the endogenous substrates Regulator of G-protein signalling RGS4 and RGS5 (22). Finally, a cross-kingdom survey revealed that N-terminal cysteinyl-dioxygenases (NCOs) are ubiquitously distributed across plants and animals, with PCO-*like* proteins also found in the chytrid lineage of fungi (21).

The most common cause of acute hypoxia for plants is flooding, due to a four orders of magnitude decrease in the gas-exchange rate in water compared to air (23). Limitation in O_2_ availability severely impacts cell metabolism, including aerobic respiration, leading to reduced ATP production (24, 25). Upon hypoxia detection, plant cells reconfigure their metabolism to ensure basal energy production even when aerobic respiration is impossible. These include the activation of fermentation and the promotion of starch and sucrose catabolism (26–28). While anaerobic metabolic adaptation is conserved across land plants, recent evidence suggests that ERFVII as N-degron substrates emerged relatively late, with the evolution of vascular plants (1). O_2_ concentrations below aerobic levels have also been reported to occur physiologically in plant tissues such as seeds, ripe fruits, and bulky storage organs (29–31). The physiological relevance of oxygen sensing is also highlighted by the developmental defects observed in the homozygous *4pco* mutant, including delayed germination and growth, increased leaf serration, and flower sterility (17, 20, 21). Recent studies also demonstrated the establishment of endogenous hypoxic niches in shoot and root meristems, thus designating the role of O_2_ as a diffusible signalling molecule (21, 32). Additionally, fluctuations in O_2_ levels between day and night have been recently reported to impact shoot growth (33).

Modulation of the O_2_ sensing pathway to anticipate, enhance or redirect regulation of adaptive genes is a highly sought goal of plant hypoxia research (34). This can be achieved by interfering with the N-degron substrates, such as the ERFVIIs, or the O_2_-dependent effectors, such as the NCOs (35). Previous studies have showed that increasing the levels or stability of ERFVII transcription factors enhances submergence and hypoxia tolerance in plants: for example, in Arabidopsis, overexpression of RAP2.2, RAP2.12, HRE1, and HRE2 improved tolerance to low-O_2_, though constitutively stable RAP2.12 impaired growth and development (5, 6, 36).

Since constitutive ERFVII stabilisation can be detrimental, precise tuning is necessary to ensure a beneficial effect. Acting on the molecular switch that regulates ERFVIIs in hypoxia, such as N-cys oxidation, could be the key to modulate intensity and dynamics of the molecular response to hypoxia. PCOs are regarded as the ideal target to manipulate, due to their more specialised activity, and thus likely reduced number of substrates, compared to ATE and PRT6/BIG. Moreover, the identification of key residues within PCO enzymes, along with examination of reported and predicted structures of NCO enzymes in which some of these amino acids are not conserved, provides an opportunity to understand how to modulate the activity of these enzymes and to predict the consequences of their molecular engineering (20, 37–39).

## Results

### Predicting hypoxia response dynamics in case of alteration of PCO properties

We first assessed the theoretical impact of varying PCO activity or levels on the expression dynamics of a generic *Hypoxia Responsive Gene* (*HRG*) by applying a previously established model based on ordinary differential equation (ODE) (40). For this prediction, we assumed a simplified cell environment with a single ERFVII (RAP2.12) and a single PCO (PCO4). We calculated the effects of varying *PCO4* expression levels, its Michaelis-Menten constants (*K_M_*) for O_2_ and RAP2.12, and its catalytic rate constant (*k_cat_*) (**Figure 1b**). Our ODE-based simulations predicted constitutively high *HRG* expression in the case of reduced PCO4 levels or activity. Alteration of substrate affinity (*K_M_*) showed high potential to enhance the magnitude of *HRG* induction under hypoxia (1% O_2_). Taken together, these predictions suggest that reducing *PCO* expression or modifying the enzyme to decrease its catalytic rate are effective strategies for priming plants against hypoxia. In contrast, altering substrate affinity is predicted to be an effective strategy to modulate the intensity of the response during hypoxia. We therefore set out to test experimentally whether NCOs with drastically different kinetic parameters would produce the effects predicted by our ODEs model, and whether this would actually help plants to cope with low-O_2_ stress.

### Sequence and structure divergence among eukaryotic NCOs

We next explored the divergence of NCO sequences within the eukaryotic domain. We selected 50 NCO sequences from the animal, plant and fungal kingdoms and reconstructed their phylogeny based on amino acid sequence comparison (**Supplementary** Figure 1a). This analysis confirmed the taxonomic clustering across the three kingdoms and guided our selection of NCOs for subsequent biochemical characterization. Specifically, we chose *Homo sapiens* ADO (HsADO) and *Amphimedon queenslandica* ADO (AqADO) from the animal kingdom, *Spizellomyces punctatus* ADO (SpADO) from fungi, and *Marchantia polymorpha* PCO (MpPCO), *Klebsormidium nitens* PCO (KnPCO), and *Arabidopsis thaliana* PCO4 (AtPCO4) from plants. HsADO was included due to its previously demonstrated ability to complement the Arabidopsis *4pco* mutant (17). AqADO was chosen as a representative of early metazoan lineage (41). SpADO belonging to the chytrid clade, represented fungal NCOs (21, 42). Finally, MpPCO and KnPCO, from bryophytes and algae, respectively, were selected to explore PCO diversity among non-plant clades that do not have ERFVIIs as Cys-degron substrates. AtPCO4 served as a control throughout the study.

We looked into structural differences among the selected NCOs. We relied on the structures which have been solved by X-ray crystallography for AtPCO4 and HsADO (PDB: 6S7E and 7REI, respectively) whereas we used the AlphaFold2-predicted structures (43) for the remaining NCOs. The core active site, comprised of two β-barrels, is conserved across all six enzymes (**Figure 2a(I)**). Analysis of the internal substrate-binding cavity revealed varying volumes: HsADO and SpADO possessed smaller pockets (276.8 Å³ and 316.8 Å³, respectively) compared to AtPCO4 (568.4 Å³). In contrast, AqADO, MpPCO, and KnPCO exhibited cavity volumes comparable to AtPCO4 (681.2 Å³, 680.6 Å³ and 587.3 Å³, respectively), suggesting a shared potential to accommodate protein N-termini (**Supplementary** Figure 2). Moreover, analysis of the electrostatic potential surrounding the active site revealed notable differences, implying variations in the interaction with substrates (**Figure 2a(II)**). For example, HsADO and AqADO exhibited a similar predicted three-dimensional structure and overall size to AtPCO4 but displayed more negative electrostatic charge around their active sites. This electrostatic difference could explain the previously demonstrated substrate preference of HsADO for positively charged peptides (22). Outside the active site, SpADO featured an additional N-terminal α-helix and an internal loop (residues 250-260) and was more positively charged than AtPCO4 at the C-terminal region. MpPCO and KnPCO showed conservation at the core active site but contained additional internal loops and a less positively charged external surface (**Figure 2a**). When zooming in on the two β-barrel domains (Domain I: residues 96-115, Domain II: residues 164-182 in AtPCO4) that coordinate the iron centre in AtPCO4 (**Figure 2b**), alignment of these domains revealed high sequence similarity, including conserved histidine residues (H98, H100, H164), which are critical for iron coordination, aspartate (D176) and tyrosine (Y182) proposed to be necessary for substrate binding in AtPCO4 (20, 39). A high level of sequence similarity for these residues was also found more generally across all the analysed NCO sequences (**Supplementary** Figure 1b).

**Figure 2.**
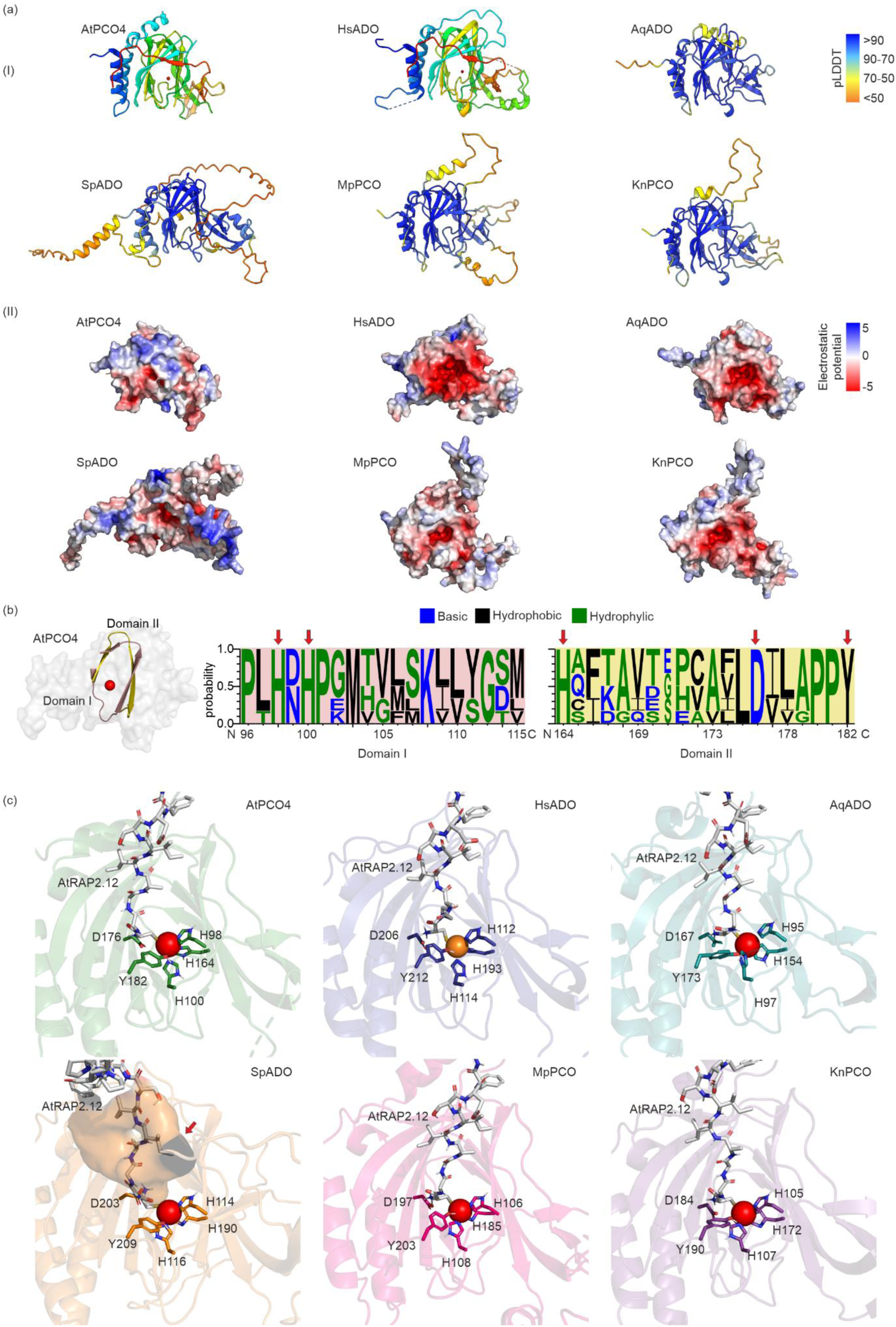
Structural comparison of selected NCOs. (**a, I**) 3D structures of AtPCO4 and HsADO (PDB: 6S7E and 7REI, respectively) and AlphaFold2 prediction of AqADO, SpADO, MpPCO and KnPCO shown in cartoon representation. AlphaFold2 predictions are coloured according to the predicted local distance difference test (pLDDT) values (red: low accuracy in prediction, blue: high accuracy in prediction). (**a, II**) Surface representation of NCO enzymes using electrostatic colour map. (**b**) Multiple sequence alignment of amino acid residues in Domain I and II of AtPCO4 active site. Amino acids are coloured by hydrophobicity. Red arrows mark ubiquitously conserved residues putatively involved in iron coordination (H98, H100, H164 in AtPCO4) and substrate binding (D176 and Y182 in AtPCO4). (**c**) Molecular docking of the first 14 amino acids of RAP2.12 (grey) inside the NCOs showed in **a**. Coloured sticks mark amino acids involved in iron coordination (His) or substrate binding (Asp and Tyr). The orange cloud in SpADO indicates the volume occupied by an extra loop which potentially interferes with substrate access.

To assess the potential for these NCOs to be functionally relevant *in vivo*, we first performed an *in silico* analysis of their interaction with the PCO-substrate RAP2.12_2-15_ peptide using molecular docking (**Figure 2c**). HsADO, AqADO, MpPCO and KnPCO exhibited similar RAP2.12_2-15_ molecular trajectories to AtPCO4. Instead, SpADO showed altered positioning, with an additional internal loop for SpADO (residues 254-259, indicated by the orange surface shading) hindering peptide access to the active site.

### NCOs vary in the catalysis ERFVII oxidation

To test the ability and efficacy of the six selected NCOs to regulate ERFVIIs in plants, we first compared their kinetic parameters for Arabidopsis RAP2.12 and oxygen. Notably, even among Arabidopsis PCOs, different isoforms exhibit different kinetic activities, with AtPCO4 having the highest catalytic turnover among the five PCOs, potentially influencing substrate preferences and different roles of PCOs under stress or physiological conditions (44). Based on this, we hypothesized that NCOs with greater sequence divergence might display even more distinct kinetic properties. While kinetic data for AtPCO4, MpPCO, and KnPCO were already available from previous studies, which demonstrated their oxygen-dependent activity on RAP2.12 (44, 45), we extended the analysis to AqADO, HsADO, and SpADO. Therefore, we expressed these three NCO proteins in *Escherichia coli* and purified them as described previously (44). We determined the kinetic parameters of AqADO and HsADO for a peptide corresponding to the first 14 amino acids of RAP2.12 (RAP2.12_2-15_). Substrate turnover using varying RAP2.12_2-15_ concentrations was used to estimate catalytic parameters via Michaelis-Menten fitting (**Figure 3a**, **Supplementary Table 1**). Subsequently, we determined the *K*_M_ (O_2_) values of AqADO and HsADO with respect to RAP2.12_2-15_ oxidation, in order to assess their oxygen-sensing potential (**Figure 3b**, **Supplementary Table 2**).

**Figure 3.**
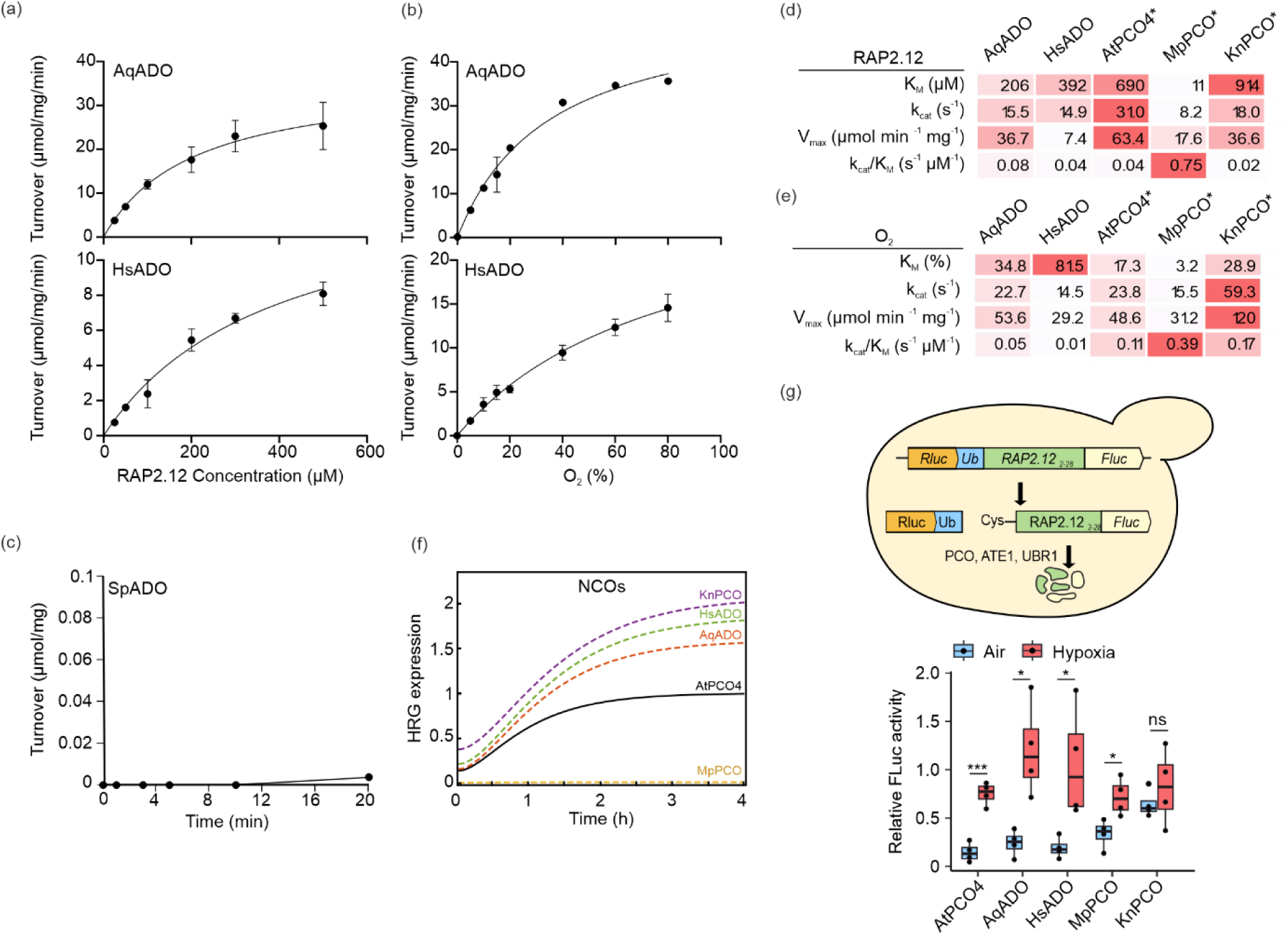
Characterization of NCO activity. Michaelis–Menten kinetic plots showing the cys dioxygenation activity of AqADO and HsADO in relation to RAP2.12_2–15_ peptide (**a**) and O₂ (**b**) concentrations. (**c**) Cys oxidation rate for SpADO using 200 µM of RAP2.12_2–15_ peptide. (**d**) Kinetic values of selected NCO towards the RAP2.12_2–15_ peptide. Colour formatting to identify highest values with strongest red hue is applied by row. (**e**) Kinetic values of selected NCO towards O_2_. Colour formatting in (**d, e**) to identify highest values with strongest red hue is applied by row. Asterisks in (**d, e**) indicate data presented in previous publications (44, 45). (**f**) ODE model prediction of *HRG* expression by RAP2.12 when this is regulated by different NCOs. (**g**) On top, a scheme depicting the DLOR assay in *S. cerevisiae* (46). On the bottom, relative FLuc activity in yeast cultures expressing NCO together with the DLOR substrate in hypoxia (1% O_2_ v/v) and normoxia (21% O_2_ v/v) for 6 hours. Asterisks indicate statistical difference between air and hypoxia as evaluated using a two-sided Student’s t-test (*n* = 4, *p* < 0.05).

It is worth noting that we used 200 µM RAP2.12 peptide in this study to determine oxygen affinity, which is lower than that used to determine kinetic parameters of AtPCO4, MpPCO and KnPCO in previous studies (44, 45).

Full kinetic data including standard errors are provided in **Supplementary Table 1**, **2**. These data confirmed that AqADO and HsADO can function as cysteine dioxygenases on the RAP2.12_2-15_ peptide in an oxygen-dependent manner, similar to AtPCO4. In contrast, SpADO showed no substrate turnover over time (**Figure 3c**), indicating its inability to oxidize RAP2.12 *in vitro* (at least under the conditions used in this study) consistent with the *in silico* predictions.

Comparative analysis of *K*_M_ values for RAP2.12_2-15_ revealed the highest peptide affinity (11 µM) for MpPCO, even higher than AtPCO4 (690 µM) or any other AtPCOs (44, 45), followed by AqADO and HsADO (206 µM and 392 µM, respectively), with KnPCO showing the lowest (914 µM) (**Figure 3d**). The high affinity of MpPCO for the RAP2.12_2-15_ peptide is particularly remarkable considering that even the most affine PCO in Arabidopsis, AtPCO2, has a *K_M_* almost 8 times higher than MpPCO (90 µM) (44).

When considering O_2_ affinity, MpPCO again showed the lowest *K*_M_ (3.2 %) (45), even lower than that reported for any other AtPCOs (44). Additionally, all the other NCOs tested in this assay displayed higher *K*_M_ (O_2_) compared to the value of 17.3 % reported for AtPCO4 in the literature, with HsADO exhibiting the lowest affinity (81.5 %). The high affinity of MpPCO for both oxygen and RAP2.12_2-15_ determined the highest catalytic efficiency (*k_cat_* / *K*_M_) (0.39 s^-1^ µM^-1^ for O_2_ and 0.75 s^-1^ µM^-1^ for RAP2.12_2-15_) among all tested NCOs, despite its low catalytic constant (*k_cat_*) (15.5 s^-1^ for O_2_ and 8.2 s^-1^ for RAP2.12_2-15_). Notably, MpPCO and KnPCO exhibited the highest and similar levels of catalytic efficiency towards O_2_ (0.39 s^-1^ µM ^-1^ and 0.17 s^-1^ µM ^-1^, respectively) (45). This suggests that MpPCO and KnPCO could be catalytically active in plant cells, and thus repress ERFVII accumulation, even at low-O_2_ concentrations.

In contrast, AqADO showed kinetic values for O_2_ similar to those of AtPCO4 but had a higher RAP2.12_2-15_ peptide affinity. This indicates that AqADO would respond similarly to AtPCO4 under decreasing O_2_ concentrations; however its strong peptide affinity suggests a stronger interaction with ERFVII substrates, which could enhance the substrate recognition and therefore NCO activity. Finally, HsADO showed comparable catalytic efficiency towards RAP2.12_2-15_ compared to AtPCO4, but reduced affinity for O_2_ (44). Although this may reflect the conditions under which this parameter was determined, it nevertheless suggests that HsADO may be unable to effectively target ERFVII in mild hypoxic conditions.

We next predicted the impact of these kinetic parameters on gene expression under hypoxia. We integrated the kinetic values of each NCO into the above-described ODE-based model to simulate generic hypoxia-responsive gene (*HRG*) expression under hypoxic conditions (1% O_2_ v/v) (**Figure 3f**). As expected, the model predicted no *HRG* induction in hypoxia with MpPCO, whereas control of RAP2.12 by AqADO, HsADO and KnPCO increased *HRG* levels compared to the levels obtained with AtPCO4.

To validate the predictions on cysteine dioxygenase activity of the selected NCOs, we utilized yeast as a heterologous expression system (**Figure 3g**). Yeast, possessing the Arg/N-degron pathway components (ATE1 and UBR1) but lacking endogenous NCOs and ERFVIIs, provides a suitable environment for assessing NCO activity without interference. We co-transformed yeast cells with each of the NCO enzymes and the Dual Luciferase Orthogonal Reporter (DLOR) system (46), enabling *in vivo* evaluation of NCO catalytic activity. Co-transformed yeast cells were subjected to hypoxia (1% O_2_ v/v) or normoxia (21% O_2_ v/v) for 6 hours. As expected, the ERFVII-based, Cys2-substrate was stabilized under hypoxia in yeast expressing AtPCO4. Significant substrate stabilization was also observed in yeast expressing AqADO, HsADO, and MpPCO. Conversely, KnPCO expression did not result in RAP2.12 accumulation under hypoxia. In fact, the substrate was already stable under normoxic conditions (**Supplementary** Figure 4). The lack of substrate stabilization in the presence of KnPCO contradicts the *in vitro* data, overall suggesting that KnPCO is unable to effectively target RAP2.12 for degradation *in vivo*.

### Substitution of endogenous PCO with different NCOs affects growth and development in Arabidopsis

To assess NCO activity *in planta* and its impact on growth and development, we first tested whether the six selected NCOs localize where known substrates are expected to be found: the cytosol and the nucleus. To this end, we transiently transformed Arabidopsis wild-type leaves with constructs bearing codon-optimized *NCO* genes, fused to GFP and placed under the control of the 35S constitutive promoter (CaMV). All selected NCOs localised to the cytosol and nucleus as expected (**Figure 4a**, **Supplementary** Figure 5).

**Figure 4.**
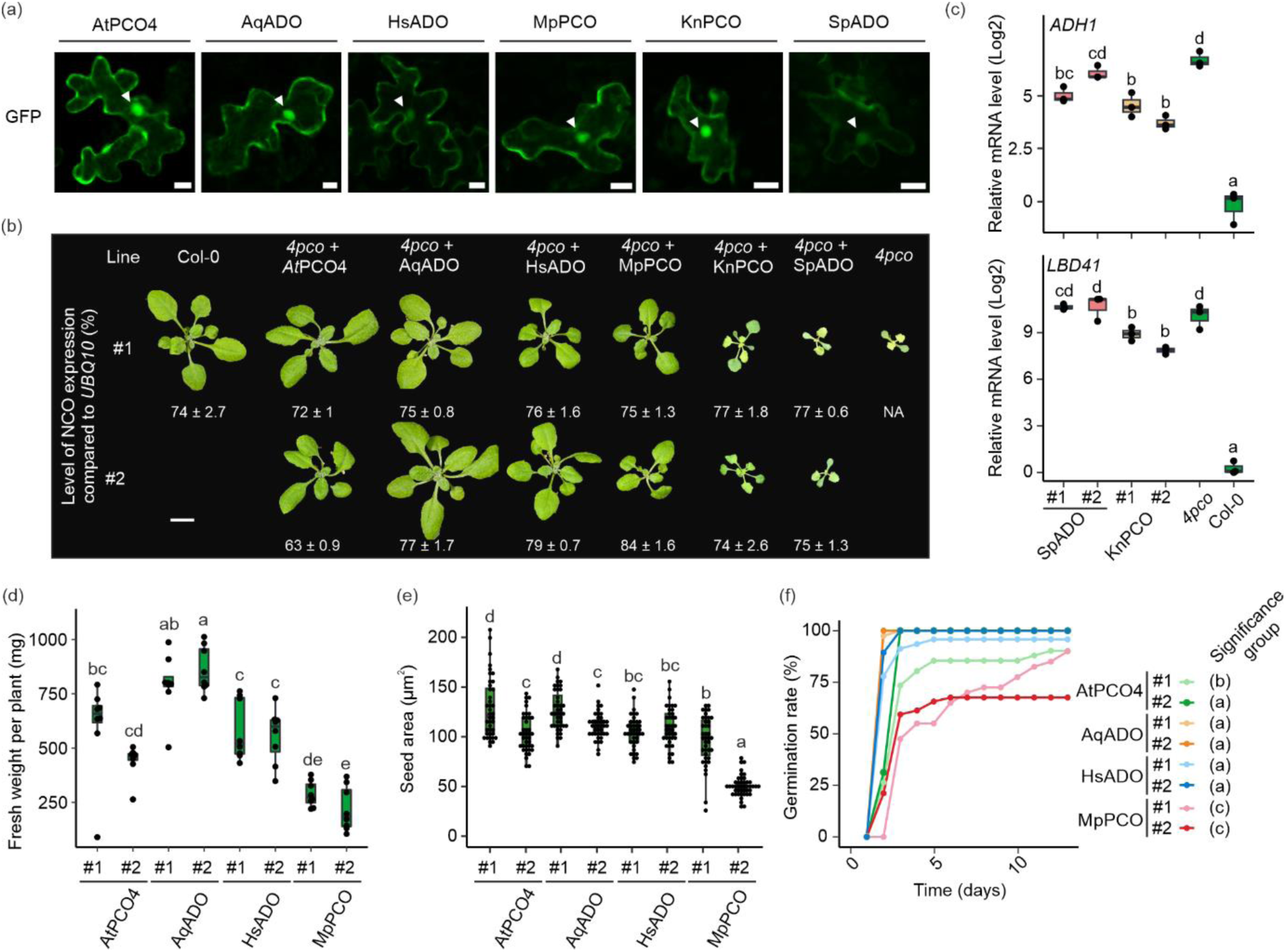
Complementation of Arabidopsis *4pco* mutant with selected NCOs. (**a**) Confocal images of NCO:GFP fusion proteins in *Agrobacterium*-infiltrated Arabidopsis leaves. Scale bar: 10 µm. (**b**) Phenotypic differences of 25-day old, *NCO*-expressing plants compared to the Col-0 wild type. Two independent lines are shown for each selected NCO. Relative Expression Level (REL) of each *NCO* compared to the expression of the *UBQ10* gene in the respective *NCO*-expressing line is shown under each plant. Scale bar: 1 cm. (**c**) Relative mRNA level (log2) of two hypoxia-responsive genes (*ADH1*, *LBD41*) in non-complementing lines expressing SpADO and KnPCO compared to Col-0 and the *4pco* mutant (*n* = 3). (**d**) Fresh weight measurement of 25-day old NCO-complemented plants (*n* ≥ 7). (**e**) Measurement of seed area of NCO-complemented lines compared to Col-0. Statistical differences were evaluated using one-way ANOVA, followed by Tukey HSD post-hoc test (*n* = 40, *p* < 0.05) (c, d, e). (**f**) Germination rate of NCO-complemented seeds after three days of stratification. Statistical difference was evaluated using the Kaplan-Meier survival analysis (*n* ≥ 49, *p* < 0.05).

To test functional complementation capacity of the six selected NCOs, we took advantage of the quadruple *4pco* mutant, which displays severe developmental and growth defects. We stably introduced the *NCO*-coding sequences in this genetic background. We placed the six *NCO*-coding genes under the control of the *AtPCO4* promoter so that they could be expressed at levels comparable with endogenous *PCOs* and subjected to the same transcriptional and post-transcriptional regulation. We selected two independent lines per NCO construct, with comparable transgene expression levels. We also verified that the insertion of the NCO-bearing T-DNA in these lines did not inactivate genes related to growth, development or O_2_ sensing (**Supplementary** Figure 6d-k).

As for PCO4 complementation, two lines with slightly different expression levels were included (**Figure 4b**) to assess the effect of expression level on plant physiology. As expected, plants expressing *AtPCO4* complemented the mutant phenotype. However, the different gene expression levels led to phenotypic differences. The lower-expressing line (#2) exhibited reduced rosette fresh weight, seed size and different germination rate compared to #1 (**Figure 4d, e, f**). Such broad phenotypic consequences related to relatively small differences in PCO4 expression, reveal the important contribution of PCOs to plant growth and fitness.

Plants expressing SpADO resembled the *4pco* mutant, including stunted growth, pale serrated leaves, and flower sterility (**Figure 4b**) (17, 21). Similarly, expression of KnPCO could not fully prevent growth defects, including pale leaves and reduced growth, although these were less severe than in *4pco* and in SpADO-complemented lines. Nonetheless, the plants could not produce seeds (**Figure 4b**). The *4pco* mutant expresses *HRG* at high levels under normoxic conditions, as a consequence of its inability to repress ERFVII activity (21). To confirm whether these lines displayed a *4pco*-like phenotype also at the transcriptional level, we assessed the expression of two *HRGs*, *ADH1* and *LBD41*, under normoxic conditions. Both *SpADO*- and *KnPCO*-expressing lines showed increased hypoxic response upon normoxia similar to *4pco* plants, indicating a constitutive hypoxic response (**Figure 4c**). These results, consistent with *in silico* and *in vitro* data (**Figure 2**, **Figure 3**), indicate that SpADO and KnPCO cannot functionally complement the *4pco* mutant.

In contrast, expression of *HsADO*, *AqADO*, and *MpPCO* rescued the *4pco* phenotype, similarly to *AtPCO4*, although we could detect differences at the level of plant vegetative growth and reproductive development among these lines (**Figure 4b**). We therefore decided to exclude non-complementing genotypes and focus our analyses on investigating the effect of complementing NCOs on plant physiology. AqADO-complemented plants showed increased fresh biomass accumulation compared to AtPCO4 #1 (**Figure 4d**). MpPCO-complemented plants exhibited decreased fresh weight, increased leaf number, delayed flowering, decreased seed area, and delayed germination, both with and without stratification, similarly to AtPCO4 #2 (**Figure 4d, e, f, Supplementary** Figure 7a-d). One of the two MpPCO-expressors, line #2, presented an additional pale seed phenotype (**Supplementary** Figure 7d). HsADO-complemented plants showed similar fresh weight and leaf number as line AtPCO4#1, although they also exhibited delayed flowering and decreased seed area (**Figure 4d, e, f, Supplementary** Figure 7a-d).

### Transcript profiling reveals a negative correlation between *HRGs* expression and growth

Next, we analysed transcriptional changes in *NCO*-expressing plants to investigate the molecular basis of the observed phenotypic differences. To this end, we sequenced the poly(A)-enriched transcriptome of 7-day-old *NCO*-expressing seedlings grown under atmospheric (normoxic) conditions. Multidimensional scaling analysis (MDS) revealed only moderate transcriptional differences between *AtPCO4*- and *AqADO*-expressing plants, while *HsADO*- and *MpPCO*-expressors exhibited more divergent transcriptomic profiles (**Supplementary** Figure 8a). Differential gene expression analysis was performed to identify genes significantly altered (FC>|1|, *p-adj* < 0.05) when compared with PCO4-complemented plants, followed by Gene Ontology (GO) enrichment analysis to assess functional differences. *MpPCO*-expressing plants showed the most up- and down-regulated genes (DEGs), with 175 upregulated and 62 downregulated genes (**Figure 5a**). AqADO-expressors exhibited mostly upregulated genes compared with AtPCO4 plants (55 up- and 13 down-regulated genes) while HsADO expression caused the least changes (25 up- and 19 down-regulated genes) (**Figure 5a**). Remarkably, the overlap between differentially expressed genes was extremely limited among NCO-expressor lines (**Figure 5b**). GO enrichment analysis revealed that expression of all three NCOs led to the up-regulation of low oxygen inducible genes, although only MpPCO-expressors exhibited higher levels of core hypoxia-response genes, especially those that require ERFVII for their up-regulation, under low oxygen (47, 48) (**Figure 5c-d, Supplementary** Figure 8c). A similar, though statistically non-significant, trend was observed in *HsADO*-expressing plants (**Figure 5d**). Additionally, *MpPCO*-expressing plants showed enrichment of GO terms associated with immune response and stress-related hormones (**Supplementary** Figure 8c). While it is difficult to explain the enhanced growth and biomass accumulation observed for *AqADO*-expressing plants based on the DEGs list, on the other hand the reduced size of *MpPCO*-expressors is potentially associated with the activation of the hypoxic response under aerobic conditions (**Figure 5c-d**), as observed in case of constitutive ERFVII stabilisation (49). These observations suggest that reduced *in planta* activity of these NCOs toward PCO substrates may lead to incomplete repression of the hypoxia response under normoxic conditions. However, the little overlap in DEGs suggests that, while HsADO and AqADO, differently from MpPCO, can effectively repress ERFVII activity, other substrates could evade the cysteine dioxygenation catalysed by them and trigger signalling events that result in alteration of gene expression.

**Figure 5.**
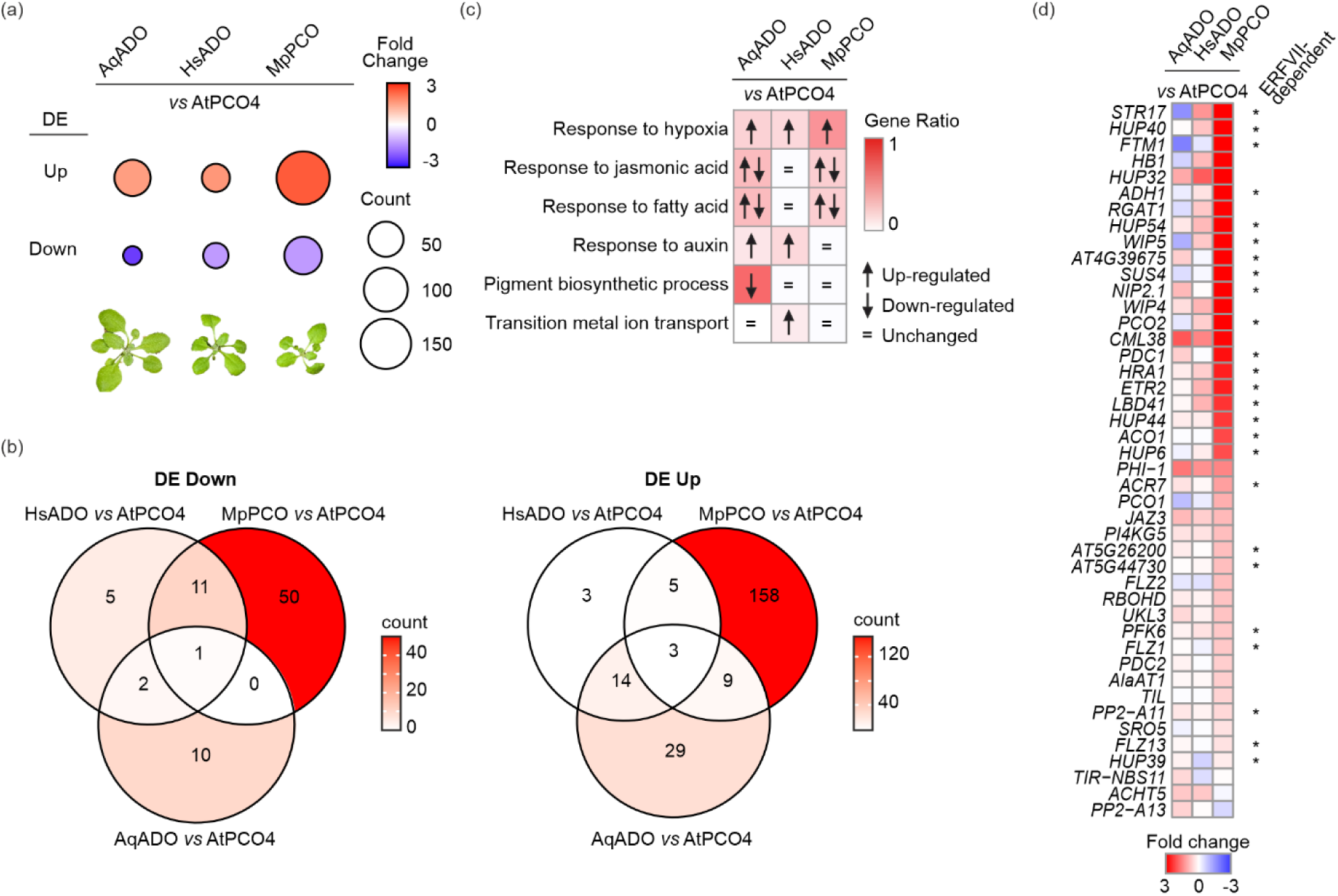
Transcriptional changes in *NCO*-expressing seedlings grown under aerobic conditions. (**a**) Differentially expressed genes in AqADO-, HsADO- and MpPCO-complemented lines compared to AtPCO4-complemented ones. (**b**) Venn diagrams showing the limited overlap of differentially upregulated and downregulated genes among the three contrasts displayed in (**a**). (**c**) Comparison of changes in main gene ontology categories among NCOs. (**d**) Differential expression of hypoxia-core genes in *NCO*-expressing lines compared to *AtPCO4*-expressing lines.

### NCOs with reduced activity on ERFVII increased tolerance to submergence

Finally, we investigated the effect of different NCO enzymes on the plant’s low-O_2_ response. First, we measured the expression of three *HRGs* (*LBD41*, *ADH1*, and *PDC1*) in plants subjected to 24 hours of normoxia (21% O_2_ v/v) or hypoxia (1% O_2_ v/v), followed by 1 hour of reoxygenation (21% O_2_ v/v) under dark conditions (**Figure 6a**). Statistical analysis revealed a significant increase in the expression of all three hypoxia-responsive genes in MpPCO #1 plants under normoxic conditions (**Figure 6a, Supplementary** Figure 9a). Also, MpPCO #2 exhibited a tendency towards elevated *HRG* expression in air, although this increase was not statistically significant. As expected, hypoxia induced expression of these genes in all genotypes. Notably, both HsADO-complemented lines and MpPCO #1 plants showed significantly higher expression levels compared to all other genotypes, including the AtPCO4 #1 control. Upon reoxygenation, *HRGs* expression decreased, though MpPCO #1 and, for *ADH1*, MpPCO #2, maintained higher expression relative to other genotypes. These findings suggest that the specific activity of each NCO enzyme can differentially modulate the hypoxic response of plants, with MpPCO leading to constitutively higher expression of hypoxia-responsive genes, which persisted during reoxygenation.

**Figure 6.**
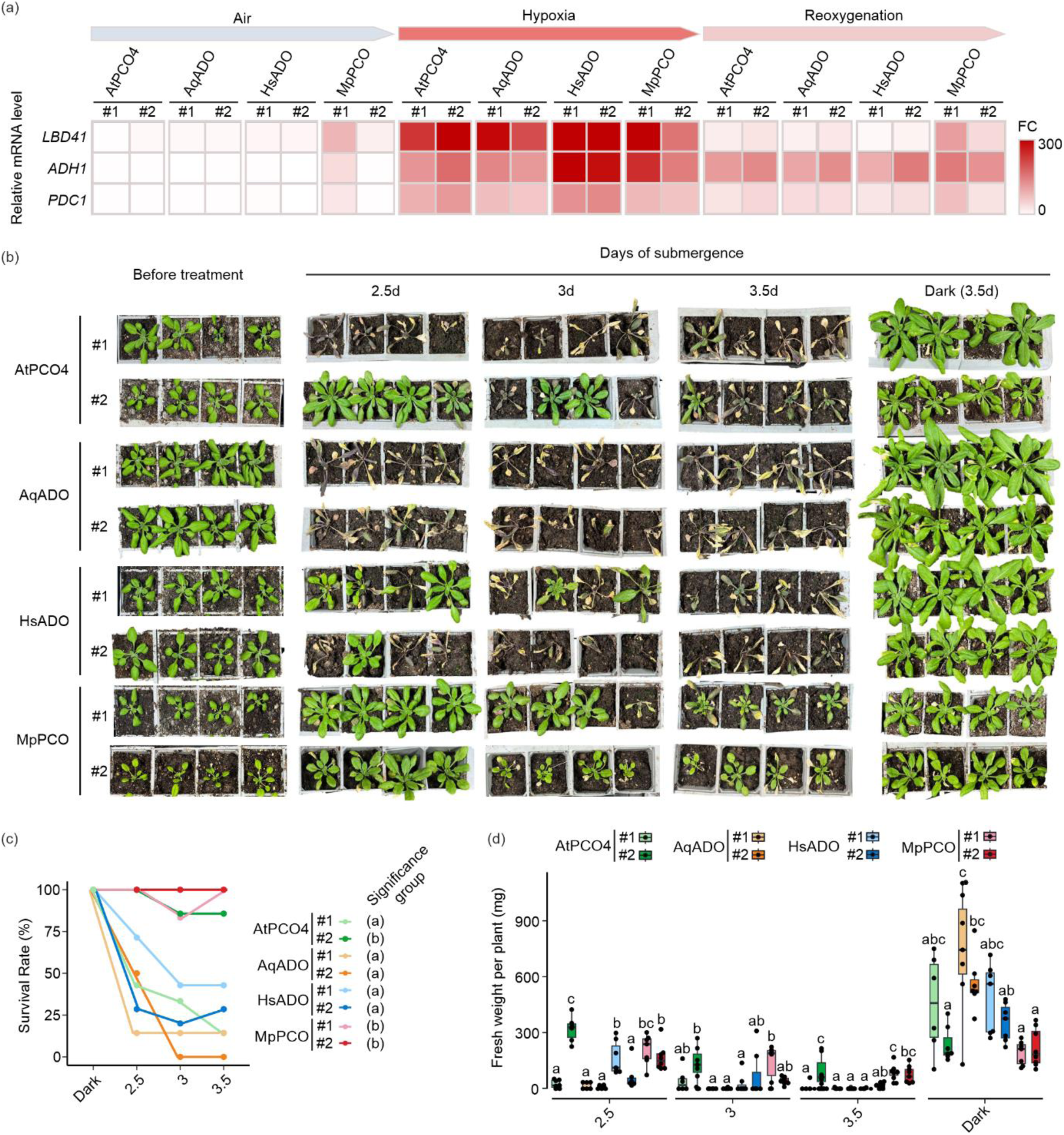
Effect of NCO-complementation under low-oxygen conditions. (**a**) Heatmap representing the relative mRNA level of three hypoxia-core responsive genes (*ADH1*, *LBD41*, *PDC1*) upon 24 hours of normoxia (21% O_2_ v/v), hypoxia (1% O_2_ v/v) or hypoxia followed by 1 hour of reoxygenation (21% O_2_ v/v). All treatments and controls were kept in the dark. Statistical significance was evaluated using one-way ANOVA among treatments, followed by Tukey HSD post-hoc test (*n* = 3, *p* < 0.05). (**b**) Phenotype of 17-day old NCO-complemented plants submerged from 2.5 to 3.5 days in the dark. Photos for each plant were taken before the treatment and four days after the end of the submergence treatment. Control plants were kept in dark for 3.5 days (*n* = 8). (**c**) Survival rate of submerged plants analysed four days after the end of the submergence treatment. Statistical analysis of survival rate was conducted using the Kaplan-Meier survival analysis method. Letters indicate significant differences among groups. (**d**) Fresh weight of plants four days after desubmergence. Statistical differences were evaluated using one-way ANOVA, followed by Tukey HSD post-hoc test (*n* ≥ 6, *p* < 0.05).

Next, we assessed the impact of NCO enzyme complementation on tolerance to low-O_2_ stress. 17-day-old soil-grown plants were subjected to full submergence in the dark for 2.5, 3, or 3.5 days. Control plants were maintained in the dark for 3.5 days. Following desubmergence and a 4-day recovery period under standard photoperiod conditions, survival rates were determined. Across all time points, *MpPCO*-expressing lines and *AtPCO4* #2 demonstrated enhanced survival (**Figure 6b, c**). HsADO-complemented lines also showed a trend of increased survival compared to AtPCO4 #1 control (**Figure 6b, c**), though to a lesser extent than MpPCO-complemented and AtPCO4 #2 lines. Already under dark conditions, these genotypes displayed slower growth and, for *MpPCO*-expressing lines, reduced biomass accumulation compared to the AtPCO4 #1 control (**Figure 6d**), consistent with the previous fresh weight and seed area measurements (**Figure 4d**). To further investigate their tolerance, we assessed tissue damage by categorizing plants as ‘dead’, ‘alive’, or ‘damaged’ (**Supplementary** Figure 9b). This analysis revealed that, despite improved survival, MpPCO- and HsADO-complemented lines still exhibited significant levels of damage compared to control conditions in the dark.

Collectively, these results indicate that plants with a more active NCO enzyme exhibited higher biomass (e.g., AqADO-complemented lines and AtPCO4 #1) and lower survival rates compared to plants with reduced NCO activity that exhibited an initial reduced fresh weight (MpPCO-complemented lines and a trend in HsADO-complemented lines), but despite experiencing damage, exhibited greater resilience to submergence.

## Discussion

In the present study, we tested the consequences of substituting endogenous Arabidopsis PCOs with homologs from different eukaryotic lineages. The motivation to undertake such a broadly cross-species approach relies on the opportunity offered by natural variation in the three eukaryotic kingdoms to explore the evolutionary space available for N-terminal cysteine oxidases. Our results confirm that variations in NCO sequences, structures and kinetic properties do impact Cys2-substrate activity, and show how this affects plant growth and development. This transgenic substitution alters the basal expression levels of genes involved in the adaptations to low oxygen availability and their dynamics when cells experience fluctuations in oxygen availability, ultimately affecting the plant’s capacity to cope with submergence conditions.

Although NCOs are ubiquitously found in plants and animals, the number of intra-specific variants varies between kingdoms. While only one or two isoform variants are found in metazoans, in flowering plants, where they constitute the main cellular oxygen sensor, their number expanded remarkably, even when considering genome duplication events (17, 21) . The observation that this is not the case for the other enzymes that participate to the N-cys-degron, such as ATE and PRT6/UBR, suggests the need, in flowering plants, for finely-tuned control of N-cys substrate stability, and the opportunity to achieve this through a cell-specific suite of PCO isoforms. For example, thorough characterisation of Arabidopsis PCOs has revealed that isoforms with limited sequence diversity exhibit rather distinct kinetic activities, potentially influencing substrate preferences and different roles of PCOs under stress or physiological conditions (44). Therefore, we speculated that NCOs with higher sequence divergence might have even more diverse kinetics.

Indeed, a comparison of six NCO from across the eukaryotic domain revealed even greater differences in affinity for the canonical ERFVII peptide and for O_2_, and activity (**Figure 3a-e**). This is most likely due to differences in sequence and structure among the enzymes, which impact, besides catalysis, substrate interaction and selectivity. Phylogenetic and structural analyses confirmed that NCOs are conserved across eukaryotic lineages, not only in their primary amino acid sequences but also in core structural features, such as the β-barrel catalytic pocket and residues essential for the iron coordination and substrate interaction (**Supplementary** Figure 1, **Figure 2**). These conserved features support a shared enzymatic mechanism and a potentially ancestral functional role. Nevertheless, we observed key structural variations among homologs, particularly in electrostatic surface charge distribution and substrate pocket volume. Such divergence likely reflects lineage-specific substrate specialization, consistent with the kingdom-specificity of known NCO targets such as plant ERFVIIs and animal RGS proteins. Our docking simulations further support this hypothesis, as the fungal SpADO displayed divergent docking trajectories for the Arabidopsis RAP2.12 peptide, suggesting substrate incompatibility. These findings also highlight the usefulness of *in silico* modelling in predicting substrate interaction, particularly for peptide-like substrates.

It is plausible that some of the tested NCOs, including ADO, may also turn over small thiol-containing molecules rather than peptide-based substrates (50). In such cases, structural features like differences in the binding pocket volume may be indicative of this substrate preference, although further biochemical validation is needed. Importantly, while structural and docking data can be informative, they do not always correlate with *in vivo* function. For instance, SpADO exhibited a markedly altered three-dimensional structure, including an additional loop potentially occluding the active site, and failed to catalyse RAP2.12 oxidation *in vitro* or complement the *4pco* mutant phenotype. Conversely, KnPCO, despite showing minimal structural deviation, was similarly inactive, suggesting potential adaptation to non-angiosperm-specific substrates or limitations in our docking approach, which focused on peptide segments rather than full proteins.

Variation in sequence and structure is also likely to impact the interaction with other proteins, including potential post-translational regulators of PCO activity. Y2H and co-immunoprecipitation surveys indicate only few potential interactions of Arabidopsis PCOs, PCO2 with the SEC5A subunit of the exocyst complex, PCO3 with the sulfurtransferase SUFE1 and PCO5 with the subunit 5A COP9 signalosome (51, 52). While this study was not designed to explore this aspect of PCO regulation, it can explain the discrepancy between the predicted and measured effects of NCOs and HRGs (**Figure 3f-g**, **Figure 6a**). The DLOR assay in yeast, where NCO expression levels are substantially comparable across cells, exhibited a trend of enhanced substrate stability in hypoxia for *HsADO*- and *AqADO-* expressing cells. However, the model indeed overestimated KnPCO and MpPCO capacity to induce Cys2-substrate degradation. This could raise doubts about the capacity of our ODE-based model to effectively predict the effect NCO substitutions on HRG expression. As stated above, protein-protein interactions are likely to impact this. Additional complications are determined by cell heterogeneity in plant tissues and the impact of NCO and ERFs transcription and translation towards the determination of *HRG* expression. Because of well-known feedback mechanisms such as *ERFVIIs* and *PCOs* gene induction by hypoxia (21, 53), *HRG* expression prediction are especially challenging in plants.

Despite the afore-described limitations, our approach defines an effective pipeline for improving the control of hypoxia responses that can be conveniently conjugated with the rational-design or directed evolution of NCO. When applied through precise gene editing (54), this has the potential to affect the plant’s ability to respond to hypoxia exactly in the desired trajectory, as the impact on gene expression in this case would be negligible. This is crucial, when looking at the phenotypes observed for plants grown under aerobic conditions in this study. Substitution of PCO4 with HsADO, AqADO, and MpPCO affected rosette biomass accumulation, flowering time, and seed size (**Figure 4, Supplementary** Figure 7). This phenotypic variability suggests that differences in kinetic efficiency or substrate interaction dynamics markedly influence growth and developmental outcomes. For example, AtPCO4 #2 line exhibited only mild *HRG* induction and yet showed marked developmental delays, consistent with computational predictions that reduced *PCO* expression drives *HRG* upregulation. Similarly, HsADO, characterized by a smaller pocket volume and reduced catalytic efficiency toward O_2_, also led to altered phenotypes in planta, including trends toward increased *HRG* expression, albeit without statistically significant differences. These results suggest that phenotypic differences likely reflect organ or tissue-specific roles for NCOs, associated with the need to regulate different substrates. The N-cys-degron substrates identified so far participate to various developmental processes, including root and leaf development and shoot apical meristem activity (3, 4, 32, 55–57). The comparison of transcript profiles of *AqADO*- and *MpPCO*-expressing plants compared to PCO4-complemented ones suggest the existence of additional substrates. There are more than 200 MC-proteins in Arabidopsis, and more Cys2-proteins can be created by endoproteolysis (6, 58).

When evaluating the effect of flooding, plants complemented with MpPCO and HsADO, which exhibited delayed seed germination, slow vegetative development, and late flowering, displayed enhanced submergence survival. These findings suggest that a trade-off may exist between growth and stress tolerance. One possibility is that enhanced hypoxic response in low-efficiency NCO-complementing lines directly influence developmental and growth regulators. However, it is also plausible that the altered phenotype is due to the regulation of other N-degron targets, such as VRN2 or ZPR2, which are known to play roles in flowering and developmental transitions. Alternatively, the link may be indirect: delayed development could allocate fewer resources to growth and more to survival mechanisms when plants experience stress. This aligns with prior studies on strategies evolved by plants to reduce growth to improve flooding tolerance due to enhanced carbon storage or metabolic flexibility (59, 60).

We consider this approach particularly valuable given the expanding roles of ERFVIIs in responding to drought, heat, salt, oxidative, and osmotic stresses, often mediated through ABA and ROS signalling pathways (61).

The elevated basal hypoxic response in MpPCO lines suggests partial stabilization of ERFVIIs under normoxia. This is supported by prior mutagenesis studies in *AtPCO4*, where disruption of substrate-binding residues (e.g., H164 and D176) resulted in increased *HRG* expression (20). While this constitutive activation is not necessarily advantageous under normal conditions, it raises the question as to whether constitutive or elevated *HRG* expression could enhance hypoxia tolerance by priming the response system. A similar concept has been observed in ethylene-treated plants, where *PHYTOGLOBIN1* induction enhances flooding tolerance (62). Similarly, pre-exposure to waterlogging has been shown to improve stress resilience via redox, transcriptional, and metabolic reprogramming (63–66). Finally, N-degron signalling has been previously linked to chromatin remodelling, via the VRN2 substrate (2, 4, 28), both of which could contribute to long-term priming to low oxygen stress. It remains an open question whether a moderate, constitutive hypoxic signature, as observed in the *MpPCO*-expressing lines, is broadly beneficial or if it risks detrimental trade-offs under non-stress conditions. These observations highlight the broader role of ERFVII signalling beyond hypoxia. NCO-mediated differential RAP2.12 stability likely affects redox homeostasis, with implications for both hypoxic stress and the subsequent reoxygenation. Modulation of ERFVIIs has been shown to impact H_2_O_2_ signalling and activate specific transcriptional responses during reoxygenation (67, 68). The phenotypic variability observed across our complemented lines likely reflects differences in energy use, redox homeostasis, and developmental state. Given this, modulating ERFVII stability via NCO activity has far-reaching implications for plant stress resilience more generally.

In summary, our findings reveal that, while the core NCO structure is conserved across eukaryotes, variations in kinetic properties, substrate pocket size and electrostatic surface potential significantly alter downstream hypoxic responses *in planta*, development and stress tolerance. This is evidenced by the differential complementation phenotypes in the Arabidopsis *4pco* mutant and the enhanced submergence tolerance observed in lines expressing less efficient *NCO* variants. These differences reflect a complex interplay between enzyme expression and kinetics, developmental regulation and ability to promptly respond to an environmental cue. By integrating structural, biochemical, and physiological analyses, this data highlights the potential for tuning the hypoxic response through targeted engineering of PCO and ultimately improve plant survival under low oxygen stress.

## Material and Methods

### Phylogenetic analysis

Phylogenetic analyses were performed using MEGA X (69). NCO sequences were aligned using the MUSCLE algorithm (70). Phylogenetic trees were obtained using the maximum-likelihood algorithm (71) and a bootstrapping method based on 1000 repeats. Multiple sequence alignment of 50 NCOs sequences and selected ones are provided in **Supplementary Table 9**, **10**.

### 3D structure prediction and molecular docking

AtPCO4 and HsADO crystal structures were retrieved from the Protein Data Bank (PDB: 6S7E and 7REI, respectively). Predicted structures for AqADO, SpADO, MpPCO and KnPCO were generated using Alphafold2 (43). Electrostatic surfaces were measured using the APBS Tool2.1 plugin in PyMOL (v3.1.3) (72). Pocket volumes were measured by the Fpocket webserver 1.0.1 (73). All structures were visualised using Pymol (74).

An AlphaFold predicted structural model of RAP2.12 (structure ID: AF-A0A1S3YXC8-F1-v4) was used to generate substrate peptide files for docking. The RAP2.12 protein was truncated to display only the first 14 amino acids (reflective of experimental conditions). The N-terminal was modified using PyMOL (v3.1.3) (74). For RAP2.12_2-15_, the N-terminal cysteine was modified to remove the hydrogen from the thiol group thereby improving metal binding (modified amino acid labelled: CYF) and the amino group was protonated. Iron-containing structures for AqADO, KnPCO, MpPCO and SpADO were obtained by alignment with AtPCO4 and subsequently complementation with Fe at the active site. Prior to docking analyses, all water molecules were removed. A 2+ charge was applied to the iron (heteroatom, HETATM) by editing the .pdb file. The protein and RAP2.12_2-15_ files were uploaded to the HADDOCK webserver (75, 76) in .pdb format. Two active residues, along with Fe which is key for coordination of the N-terminal Cys, were selected for each NCO: AtPCO4 (D164, Y182), HsADO (D206, Y212), AqADO (D167, Y173), SpADO (D203, Y209), MpPCO (D197, Y203), KnPCO (D184, Y190). HADDOCK was executed using default input parameters. Three stages of docking were applied: (1) randomisation of orientations and rigid-body minimisation (it0) (2) semi-flexible simulated annealing in torsion angle space (3) refinement in Cartesian space with explicit solvent (water), to generate 1000 models, best 200 of which were refined in stages 2 and 3. The final docked models were clustered based on similarities (RMSD) and assigned a z-score indicating how many standard deviations from the average the cluster is located. pdb files of docked results were visualised in PyMOL (v 3.1.3) (74).

### Modelling *HRG* expression as a function of NCO parameters

To describe the expression dynamics of Hypoxia Responsive Genes (*HRGs*) in the N-degron pathway-based O_2_ sensing system, we employed a previously developed mathematical model which consists of two coupled Ordinary Differential Equations (ODEs) (40).

For simplification, we assumed that all reactions occur in a homogeneous environment, disregarding compartmentalization or microenvironments within the cell. Additionally, we assumed that all reaction species – including RAP2.12, AtPCO4, O_2_, and mRNA – are uniformly distributed throughout the system. To ensure comparability, we set the initial concentration of AtPCO4 at steady state under normoxic conditions, representing the baseline (0 timepoint) in our simulations. Furthermore, we assumed that AtPCO4 protein levels remain constant following exposure to hypoxia (1% O_2_ v/v). This assumption is reasonable within a short timeframe, as protein synthesis is significantly reduced under hypoxic conditions (77). By keeping AtPCO4 levels unchanged, we can analyse the dynamic hypoxic responses based solely on its catalytic activity and affinities for O_2_ and RAP2.12.

For our simulations, we used previously fitted rate constants – *k_3_*, *K_M_*_1_, and *K_M_*_2_ – representing AtPCO4 catalytic activity and its affinities for O_2_ and RAP2.12 in the AtPCO4-mediated degradation of RAP2.12 (40). To model hypoxic responses, we obtained the initial concentrations of RAP2.12 and HRG mRNA from steady-state simulations performed under normoxic conditions (21% O_2_ v/v) for 6 hours. Subsequently, we conducted dynamic simulations under 1 % O_2_ v/v conditions for 4 hours. To explore how the N-degron pathway responds when regulated by enzymes with different kinetic properties than AtPCO4, we varied *k_3_*, *K_M_*_1_, and *K_M_*_2_ across a range of 10% to 300%, mimicking different catalytic activities and affinities for O_2_ and RAP2.12. This approach allowed us to assess how the dynamics of hypoxic responses vary when regulated by NCO from different species through the N-degron pathway.

For additional *HRG* simulations based on different NCOs, the parameters were obtained by using MATLAB’s *‘Talwar’*, *‘Fair’* and *‘Huber’* weight functions within the *nlinfit* function. The rate constants k_6_, K_M4_, and K_M5_ as well as k_9_, K_M10_, and K_M11_ were fitted for the AqADO- and HsADO-mediated degradation of RAP2.12, respectively. Meanwhile, k_7_, K_M6_, and K_M7_, and k_8_, K_M8_, and K_M9_ were re-fitted for the MpPCO- and KnPCO-mediated degradation of RAP2.12 (45). This three-dimensional nonlinear fitting is depicted in **Supplementary** Figure 3. All parameters and equations are shown in **Supplementary Tables 3 - 5.**

### Plant Material and Growth Condition

*A. thaliana* Columbia-0 (Col-0) was used as wild-type ecotype. Quadruple mutant *4pco* seeds were obtained as previously described (17). Seeds were sown in a soil:vermiculite 3:1 ratio mixture, stratified at 4°C in the dark for three days, then germinated at 22°C/20°C with a 16h:8h, light:dark photoperiod and 100 μmol photons m^-2^ s^-1^ intensity. For *in vitro* propagation, sterilized seeds were cultivated on half-strength MS (78) (Duchefa) medium, supplemented with 1% (w/v) sucrose and 0.8% (w/v) agar, and grown at 22°C/20°C,16:8, day:night photoperiod and 100 μmol photons m^-2^ s^-1^ intensity, after stratification at 4°C for three days in the dark. For leaf agroinfiltration, Col-0 plants were grown at 22°C,12:12 day:night photoperiod for three weeks and subsequently used for *Agrobacterium* infiltration as previously described (79).

### Construct design and plant transformation

For PCO substitution, cDNAs codon-optimized sequences for *Homo sapiens ADO* (*HsADO*), *Amphimedon queenslandica* ADO (*AqADO*), *Spizellomyces punctatus* ADO (*SpADO*), *Marchantia polymorpha* PCO (*MpPCO*), *Klebsormidium nitens* PCO (*KnPCO*)*, Arabidopsis thaliana PCO4* (*AtPCO4*) were synthetized (GeneArt, Thermo Fisher Scientific) and cloned into pENTR™/D-TOPO™ (Thermo Fisher Scientific™). pENTR-*HsADO*, pENTR-*MpPCO*, pENTR-*KnPCO*, pENTR-*AqADO* and pENTR-*SpADO* were recombined in pH7pPCO4GW (20) using the Gateway™ LR Clonase™ II Enzyme mix (Thermo Fisher Scientific™).

The recombined vectors were used to obtain stable transgenic lines in Col-0 using *Agrobacterium*-mediated transformation using the floral dip medium as previously described (80). Seeds were harvested and surface sterilized with 70% ethanol and 10% commercial bleach, followed by six washes in sterile water. T_0_ seeds were screened for hygromycin resistance and subsequently transferred in soil. The presence of the transgene and the T-DNA inside the sequence of the *PCO5* gene, to verify its homozygous/heterozygous state, was detected by PCR using PCRBIO Taq Mix Red polymerase (PCR Biosystems) (**Supplementary** Figure 6a-c). The sequences of primers used for screening transgene in **Supplementary Table 6**. T_3_ transgenic plants were used for experimental procedures.

For *Agrobacterium*-mediated transient expression in Col-0 leaves, NCO entry vectors were recombined in pK7WGF2 gateway.

### TAIL-seq based identification of T-DNA insertions

Thermal asymmetric interlaced polymerase chain reaction (TAIL-PCR) was used to identify the DNA flanking regions to the NCO transgene insertion. High quality DNA was extracted for NCO-complemented lines using the Monarch® Genomic DNA Purification Kit (New England Biolabs). Rounds of TAIL-PCR were conducted as previously described (81) with insertion-specific primers listed in **Supplementary Table 7**. TAIL-PCR products were purified and sequenced using the Nanopore platform (Source Bioscience Ltd). Raw reads were aligned to the *A. thaliana* TAIR10 genome with *minimap2* (v2.24) (82) using -ax map-ont -A2 -B4 -O6,24 -E2,1 -k9 -w2, and insertion sites were identified with *SAMtools* (v1.16) (83).

### Measurement of physiological traits

To analyse phenotypical differences, 17-day old plants grown in soil were used to measure fresh weight and leaf number, using at least five biological replicates.

### Confocal imaging

Agroinfiltrated Arabidopsis leaf discs were used for GFP localization. For nuclear localization, leaf discs were stained with 1 µg mL^-1^ DAPI (Invitrogen™) for 20 minutes, followed by three washes in PBS before visualization. Imaging was performed using ZEISS LSM 880 Airyscan microscope, equipped with a 20x objective lens, upon laser excitation at 405 nm and collection at 410-492 nm, for DAPI excitation, or at 488 nm and collection at 495-560 nm, for GFP imaging.

### Seed area and germination rate

For determining seed area, 40 dried seeds were imaged using ZEISS Stemi 508 stereo microscope and then measured using the ImageJ software (84).

For germination assays, eight seeds per genotype were sowed in the same plate of MS medium supplemented with 1% sucrose with or without 3 days of stratification at 4°C. Five biological replicates were analysed. The germination rates were recorded every 24 hours for 13 days. Germination was considered complete at the first protrusion of the radical.

### RNA extraction and RT-qPCR analysis

Total RNA was extracted from 60-80 mg of plant material using the phenol-chloroform extraction method as described previously (8). One microgram of total RNA was subjected to DNase Treatment (Thermo Scientific™) and retro-transcribed using Maxima Reverse Transcriptase (Thermo Scientific™). Real-time quantitative PCR was performed with QuantStudio 5™ Real-Time PCR System (Applied Biosystems) using a SYBR® Green PCR Master Mix (Thermo Fisher Scientific™). *Ubiquitn10* (AT4G0532) was used as housekeeping gene. Target genes relative expression level was analysed using the comparative 2^−ΔΔCt^ method (85). Three biological replicates were extracted for each condition, each represented by two technical replicates and the average expression was calculated. The sequences of primers used for cDNA amplification is listed in **Supplementary Table 8**.

### RNA-sequencing

*NCO*-expressing seeds were stratified for one week and subsequently grown vertically on ½ MS medium supplemented with 1% sucrose under a 16:8 light/dark photoperiod for 7 days. Samples were harvested at the emergence of the first true leaves and immediately frozen in liquid nitrogen. Total RNA was extracted using the GeneJET RNA Purification Kit (Thermo Scientific™), following the manufacturer’s instructions. RNA sequencing was performed in paired-end mode (PE150) on the NovaSeq 6000 platform (Illumina) by Novogene. Transcriptomic analyses were conducted using R software (v4.3.1). After quality assessment with FastQC, reads were aligned to the *Arabidopsis thaliana* reference genome (TAIR10) using the *Rsubread* package (v2.16.1) (86) and quantified using *featureCounts* package (87). A Multi-Dimensional Scaling (MDS) plot was generated to evaluate similarities and differences in gene expression profiles across samples. Differentially expressed genes (DEGs) were identified using the *edgeR* package (v3.42.4) (88). Fold change data and TPM are provided in **Supplementary Tables 11, 12**. Genes with an absolute log2 fold change ≥ 1 and a false discovery rate (FDR) < 0.05 (**Supplementary Table 11**) were selected for further analysis. Gene Ontology (GO) term enrichment analysis of the DEGs was performed using the *clusterProfiler* package (v4.10.1) (89). Complete GO results are provided in **Supplementary Tables 13-15.**

### Low-O_2_ treatment

NCO-complemented seedlings were grown in vertical plates for one week and subsequently treated with normoxic (21% O_2_ v/v) for hypoxic (1% O_2_ v/v) conditions for 24 hours using the H35 Hypoxistation (Don Whitley Scientific). During both treatments, plates were kept at dark to avoid O_2_ release from photosynthesis. For reoxygenation, after the hypoxic treatment plates were then exposed to normoxic condition for 1 hour, at dark. For each condition, 3 biological replicates were employed, each consisting of 10 seedlings.

### Submergence treatment

For submergence treatment, seeds were sown in a Sinclair Modular Seed soil:John Innes compost No. 2:vermiculite 2:2:1 ratio mixture, stratified at 4°C in the dark for three days, then germinated at 22°C/20°C with a 16h:8h, light:dark photoperiod and 100 μmol photons m^-2^ s^-1^ intensity. After 17 days, plants were submerged with approximately 23 cm of tap water in black tanks for 2.5 to 3.5 days, with control plants maintained at dark only for the longest duration of the treatment. After the treatment, plants were moved to aerobic photoperiodic growth condition. Survival was measured four days after desubmergence, and plants were scored as ‘alive’ for plants that produced new leaves, ‘damaged’ for plants with at least one dead leaf, and ‘dead’ otherwise. Fresh weight was measured four days after the end of the treatment. Survival analysis was conducted using the Kaplan-Meyer test.

### Luciferase measurements

Total proteins were extracted in passive lysis buffer (Promega). Firefly Luciferase activities were measured using the ONE-Glo Luciferase Assay kit (Promega) and the Nano-Glo® Luciferase Assay System (Promega) was used to measure the activity of NanoLuc® luciferase enzyme. Luciferase signal was normalized based on total protein concentration using Bradford assay (90).

### Heterologous NCO proteins expression and purification

The coding sequences of AqADO and SpADO were cloned into the NdeI and XhoI restriction sites of pET28a (Novagen) as described in White et al. 2018 (44) before transformation into *E. coli* BL21 (DE3) (New England Biolabs) for protein production. Expression of the heterologous protein and purification was also carried out as described in White et al. 2018 (44).

### Enzyme kinetic characterisation of AqADO, HsADO and SpADO

The kinetic assays were performed according to the protocol with slight modifications (91). To determine the affinities of enzymes for RAP2.12, the previously synthesized 14-amino-acid RAP2.12_2–15_ peptide (GL Biochem) (44) was prepared in reaction buffer at pH 8.0 (50 mM bis-tris propane [1,3-bis(tris(hydroxymethyl)methylamino)propane] and 50 mM NaCl dissolved in LC/MS-grade water). To maintain a reductive environment, 5 mM TCEP, 20 µM FeSO₄, and 1 mM ascorbate were added to the samples. Aliquots (100 µL) of RAP2.12_2–15_ at different concentrations (25, 50, 100, 200, 300, and 500 µM) were transferred into silicone-sealed MS vials and equilibrated with an 80:20 nitrogen-to-oxygen ratio using a mass flow controller (Brooks Instruments) to mimic atmospheric conditions. The samples were then pre-incubated in a benchtop thermocycler (Eppendorf) at 25°C for 10 min. Reactions were initiated by injecting 1 µL of enzyme at a final concentration of 0.1 µM using a gas-tight syringe (Hamilton) and were allowed to proceed for 2 min (within the linear range of activity). Reactions were terminated by injecting 100 µL of 10% formic acid, and if necessary, samples were further diluted in 5% formic acid to prevent detector saturation. The same method was applied to determine enzyme affinities for O_2_, except that 200 µM aliquots of RAP2.12_2–15_ were equilibrated with different nitrogen-to-oxygen ratios to mimic O₂ levels of 0%, 5%, 10%, 15%, 20%, 40%, 60%, and 80%. Oxidation was monitored using ultrahigh-performance liquid chromatography-mass spectrometry (UPLC-MS), and turnover was quantified by comparing the areas under the product and substrate ion peaks extracted from the total ion current chromatogram. UPLC-MS measurements were obtained as previously described (44). All kinetic figures and parameters were generated using Prism (GraphPad).

### DLOR assay using *S. cerevisiae*

A haploid parental strain BY4742 (Matα; his3-Δ1; leu2-Δ0; lys2-Δ0; ura3-Δ0) was co-transformed following the LiAc/SS carrier DNA/PEG method (92) with the DLOR-pAG413GPD (35) and each pENTR-*NCO* recombined into pAG415GPD (Addgene plasmid #14146) via LR. Transformants were selected on yeast synthetic drop-out medium lacking leucine and histidine; 6.7 g L-1 Yeast Nitrogen Base (DIFCO), 1.37 g L-1 Yeast Dropout Medium (Sigma-Aldrich) and 20 g L-1 glucose, plus 0.16 M uracil and 0.32 M tryptophan. Positive colonies were grown in a 96-well plate, as described previously (35). For hypoxic treatments, cells were incubated in the H35 Hypoxistation (Don Whitley Scientific), at 1% O_2_ (v/v), 30°C for 6 hours. Luciferase activity was measured using the Dual-Luciferase® Reporter (DLR™) Assay System (Promega).

### Statistical Analyses

All the statistical analyses were performed using R Statistical Software (v4.4.2, Foundation for Statistical Computing, Vienna, Austria). Additional details are provided in the corresponding sections.

### Additional Software

Graphs were made using and R Statistical Software (v4.4.2, Foundation for Statistical Computing, Vienna, Austria).

## Data availability

RNA sequencing raw data generated for this study was deposited in the Sequence Read Archive (SRA) at the National Centre for Biotechnology Information under BioProject ID PRJNA1292127.

## Acknowledgements

This work was supported by the European Research Council (ERC Consolidator grant 101001320); the Italian Ministry of University and research (MIUR, PRIN grant no. 20173EWRT9); and the Biotechnology and Biological Sciences Research Council (BBSRC) (BB/X001059/1) to F.L. E.F. was supported by the ERC Consolidator Grant 864888.The BBSRC grant BB/Y512953/1 supported M.L-P. A.P. was funded in part by the Engineering and Physical Sciences Research Council under projects EP/Y014073/1, EP/X031470/1 and APP77999. M.P. and Y. H. were supported by Biotechnology and Biological Sciences Research Council (UKRI-BBSRC) funding [grant number BB/T008784/1]. M.P. and Y. H. were also supported by the Clarendon Scholarship at St Edmund Hall and the Lorna Casselton Memorial Scholarship at St Cross College.

## Author Contributions

M.P. and F.L. conceived and designed the research. M.P., Y. H., X.C, M. L-P., R. L. and T.B. conducted experiments and analysed the data. Y.H. designed and applied mathematical modelling under the supervision of A.P., M.P. and F.L. wrote the manuscript with input from all authors.

