## Supplementary Figures and Tables for "Manipulating plant oxygen sensing through NCO substitution reveals trade-offs between growth and flooding tolerance"

**Supplementary Figure 1 – Phylogenetic analysis of NCO sequences.** (**a**) Evolutionary conservation of 50 NCO sequences across the three eukaryotic kingdoms (plants in green, fungi in orange, animals in blue), inferred by using the Maximum Likelihood method and JTT matrix-based model. The size of the circles on branches indicates the bootstrap values (1000 bootstrap replicates). (**b**) Conservation of amino acidic residues constituting the AtPCO4 active site in 50 NCO sequences. Amino acids are coloured by kingdom affiliation and their position in AtPCO4 is shown on the bottom.

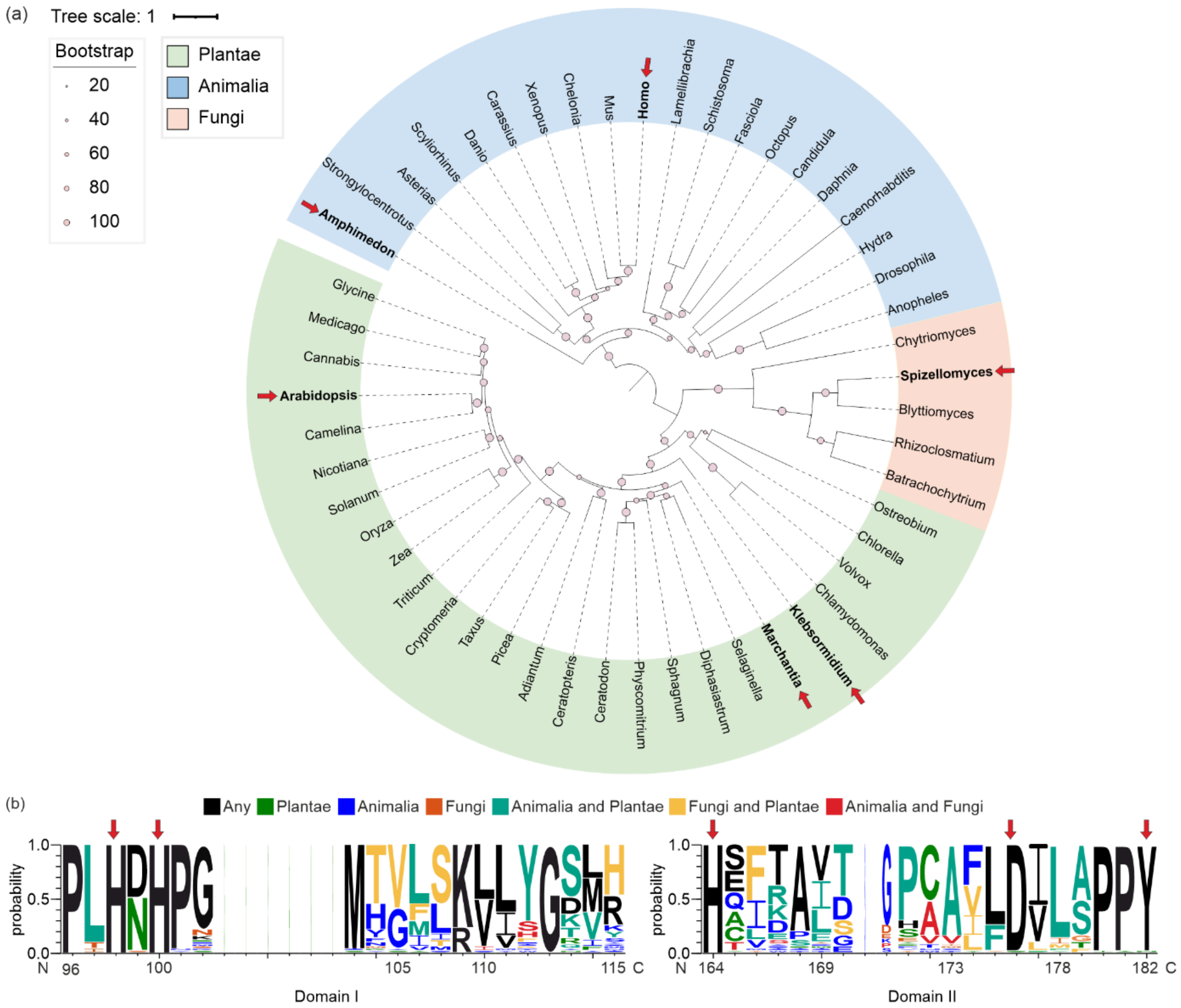

**Supplementary Figure 2 – Predicted internal pockets in selected NCOs**. (**a**) Structural differences among putative substrate-binding pockets (shown in surface representation) in 3D structures of AtPCO4 and HsADO (PDB: 6S7E and 7REI, respectively) and AlphaFold2 prediction of AqADO, SpADO, MpPCO and KnPCO (shown in cartoon representation). (**b**) Binding pocket volume (Å$^3$) and residues surrounding the pocket region. Substrate pockets were identified using the FPocket webserver (73).

(a)

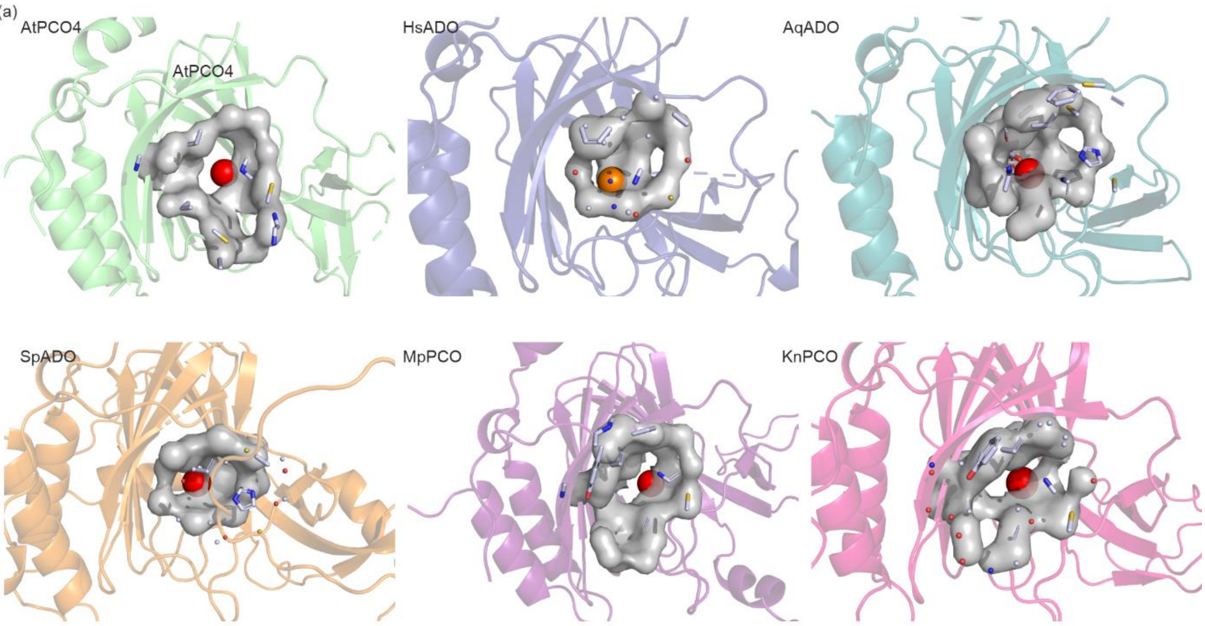


(b)

| | AtPCO4 | HsADO | AqADO | SpADO | MpPCO | KnPCO |
|---|---|---|---|---|---|---|
| Pocket Volume (Å³) | 568.4 | 276.8 | 681.2 | 316.8 | 680.6 | 587.3 |
| Atoms involved | Y73, S83, G85, F87, I95, H98,N99, H199, M103, D176, L188, P181, Y182, S183, H189, C190, Y191, F227, I228 | F101, I109, H112, H114, H193, I195, F204, D206, L208, Y212, C220, Y222, F256, C258 | F70, N72, E75, N80, A82, F84, M92, H95, H97, M104, H154, I156, F165, D166, L169, Y173, C180, Y183, F209, C211 | F70, N72, E75, N80, A82, F84, M92, H95, H97, M104, H154, I156, F165, D167, L169, Y173, C181, Y183, F209, C211 | Y81, E86, S91, G93, F95, I103, H106, H108, F187, D197, L199, P202, Y203, C211, Y213, F252, V254, Q255, R256 | Y80, E85, S90, G92, F94, I102, H105, H107, F174, D184, V186, Y190, C198, Y200, I238 |

**Supplementary Figure 3 – 3D Visualization and non-linear kinetic fitting of RAP2.12/NCO systems.** The 3D surface plot (grey) illustrates the predicted activity of NCOs including AqADO (**a**), HsADO (**b**), MpPCO (**c**) and KnPCO (**d**), as a function of both oxygen and RAP2.12 concentrations. Red curves represent enzyme kinetics across varying substrate concentrations, while blue curves depict kinetics across different oxygen levels. Experimental kinetic data for KnPCO and MpPCO were extracted from Taylor-Kearney, *et al.,* 2022 (45).

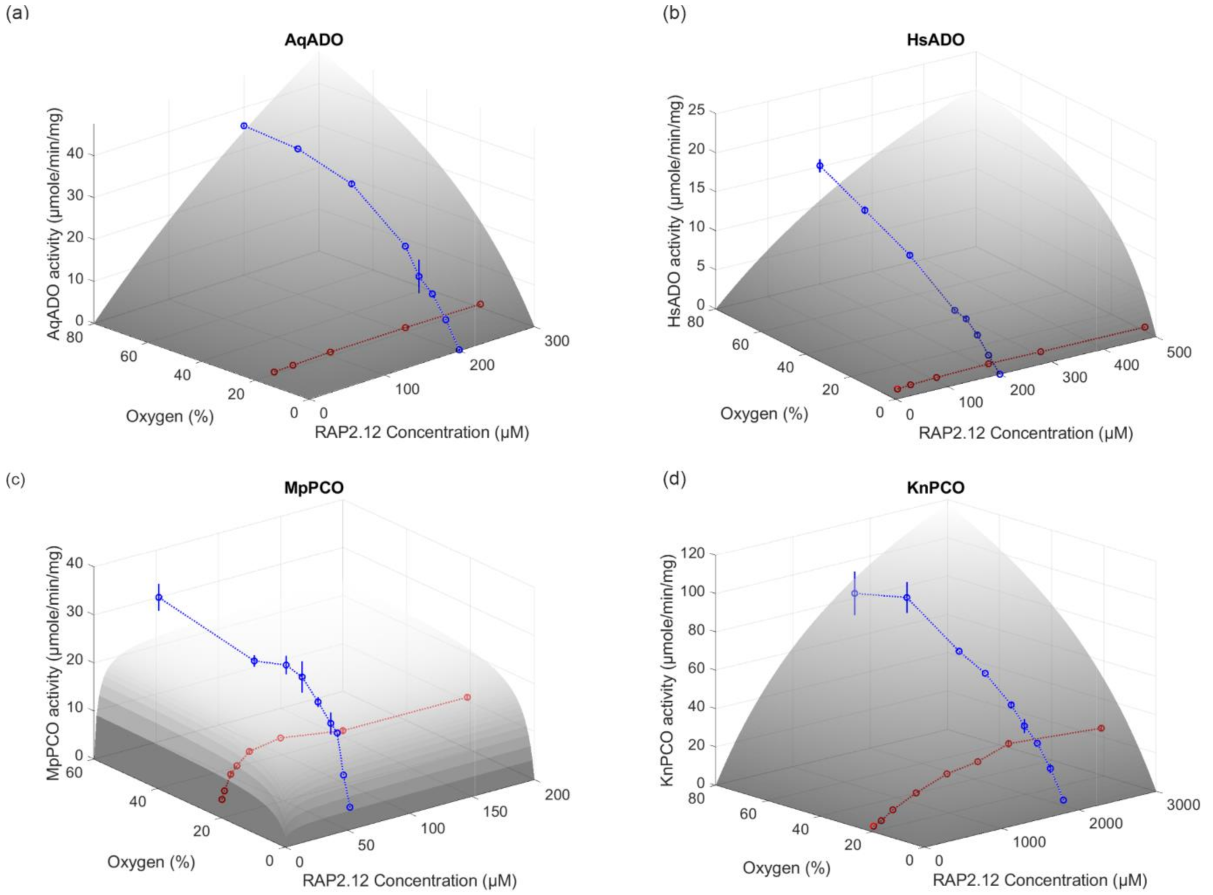

**Supplementary Figure 4 – Differential DLOR substrate stabilization in NCO-expressing yeast.** Relative FLuc activity in yeast cultures expressing NCO together with the DLOR substrate in normoxia (21% $O_2$ v/v) (**a**) or hypoxia (1% $O_2$ v/v) (**b**) for 6 hours. Letters indicate statistical difference between groups as evaluated using one-way ANOVA, followed by Tukey HSD post-hoc test ($n$ = 4, $p$ < 0.05).

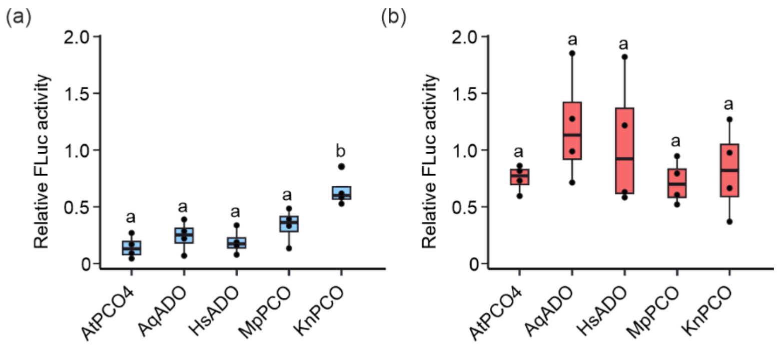

**Supplementary Figure 5 – Nuclear localization of NCO proteins.** Confocal images of NCO:GFP fusion proteins in *Agrobacterium*-infiltrated leaves. Localization in the nuclei was confirmed by DAPI staining. From top to bottom: GFP, chlorophyll, DAPI and overlay channels. White arrow indicates the nuclei. Scale bar: 10 μm.

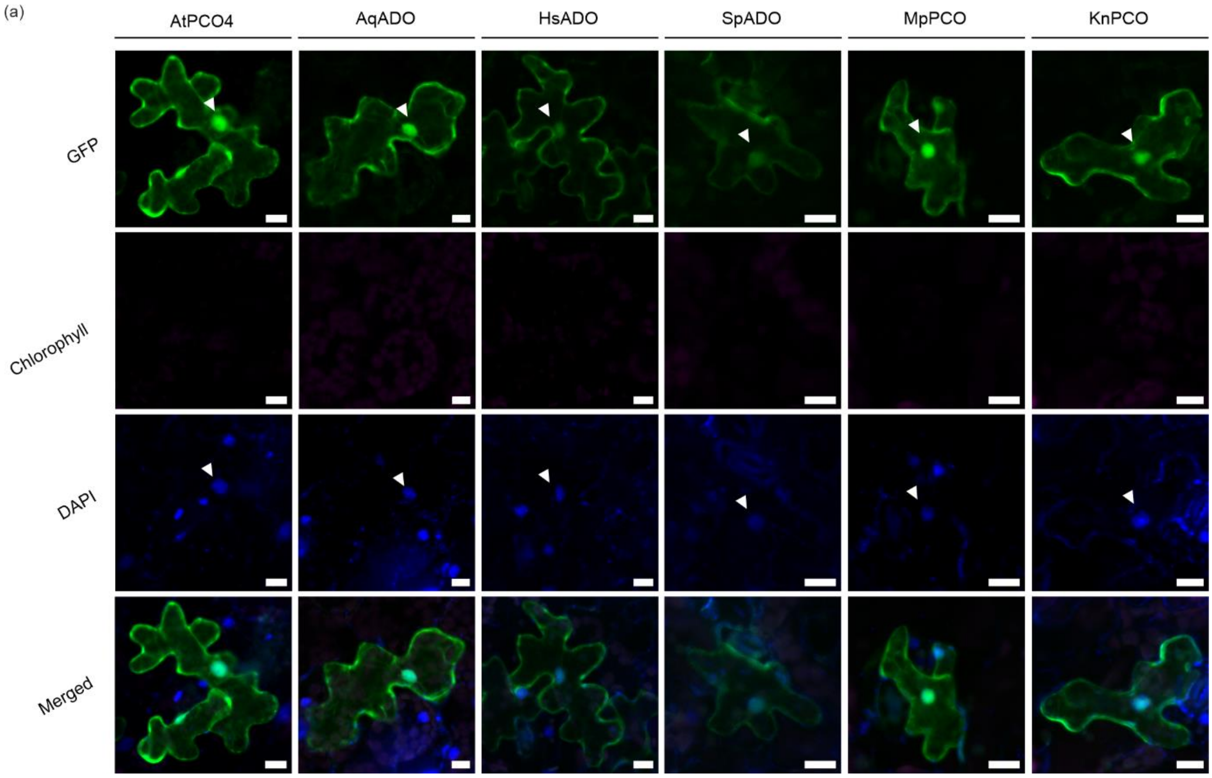

**Supplementary Figure 6 – Genotyping of NCO-complemented plants.** PCR detection of NCO-complemented plants in *4pco* homozygous background. (**a**) PCR detection of NCO-specific transgene in each independent line. The vector used to transform *Agrobacterium* was used as a positive control; Col-0 DNA and $H_2O$ were used as negative controls. (**b**) PCR detection of T-DNA in *PCO5*. DNA from *4pco* homozygous plants was used as a positive control; Col-0 DNA and $H_2O$ were used as negative controls. (**c**) PCR detection of *PCO5*. DNA from Col-0 plants was used as a positive control, DNA from *4pco* homozygous plants and $H_2O$ were used as negative controls. M: Marker. (**d-k**) Genomic location of NCO transgene insertion in NCO-complementing lines identified via TAIL-PCR. Red arrows indicate the position of the T-DNA insertion.

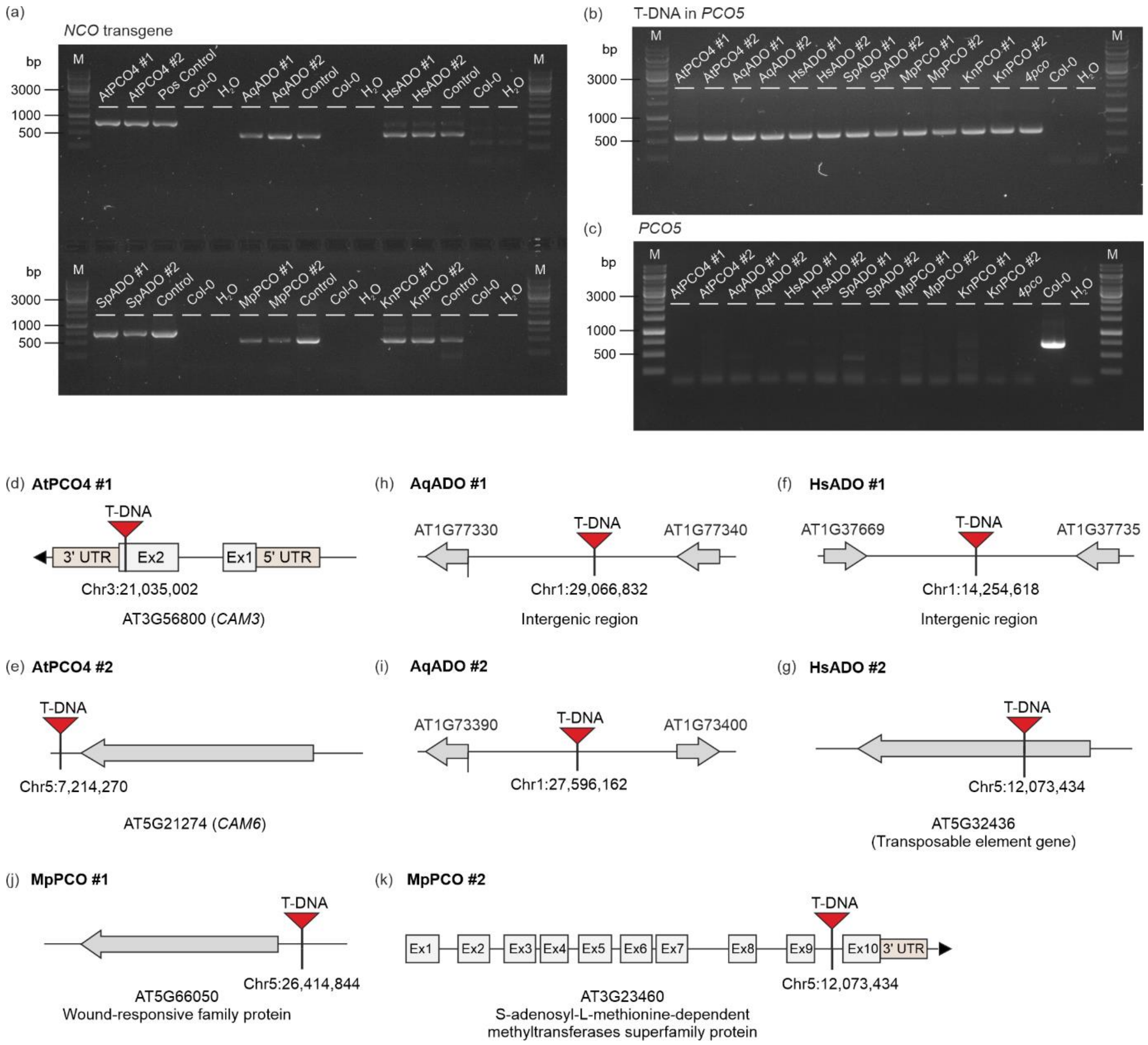

**Supplementary Figure 7 – Phenotypic effect of complementation of Arabidopsis *4pco* mutant with selected NCOs.** (**a**) Measurement of leaf number in 17-day old NCO-complemented seedlings. Statistical differences were evaluated using one-way ANOVA, followed by Tukey HSD post-hoc test ($n = 5$, $p < 0.05$). (**b**) Differences in flowering time among selected 25-day old NCO-complemented plant ($n = 5$). (**c**) Germination rate of NCO-complemented seeds without prior stratification. Statistical difference was evaluated using the Kaplan-Meier survival analysis ($n \geq 49$, $p < 0.05$). (**d**) Phenotype of seeds in different NCO-complemented plants compared to the Col-0 wild type. Scale bar: 0.2 mm.

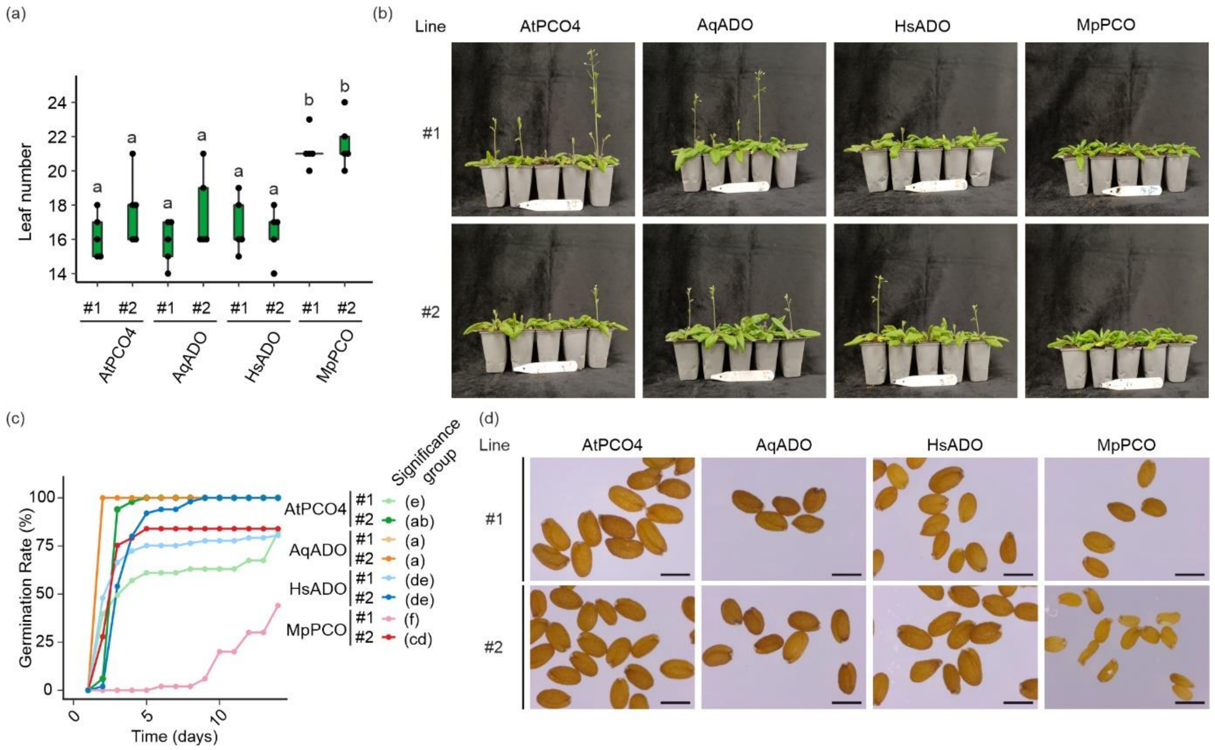

**Supplementary Figure 8 – Transcriptional changes in *NCO*-expressing seedlings.** (**a**) Multidimensional scaling plot (MDS) conducted on the normalized gene expression values of 7-day old *NCO*-expressing seedlings. Horizontal and vertical coordinates show MDS1 and MDS2, respectively, with the amount of variance contained in each component (22% and 21%). Each point in the plot represents a biological replicate. Two independent lines with two biological replicates each were sequences per NCO-complementation. Differences in colour and symbols indicate different NCO-complementation and independent lines, respectively. (**b-c**) Pairwise analysis of GO terms associated with transcripts differentially down-regulated (**b**) and up-regulated (**c**) in *AqADO*-, *HsADO*- and *MpPCO*-expressing lines compared to AtPCO4. Circle size indicates the gene ratio per GO term, with color maps indicating the False Discovery Rate (FDR) value (*p.adjust*).

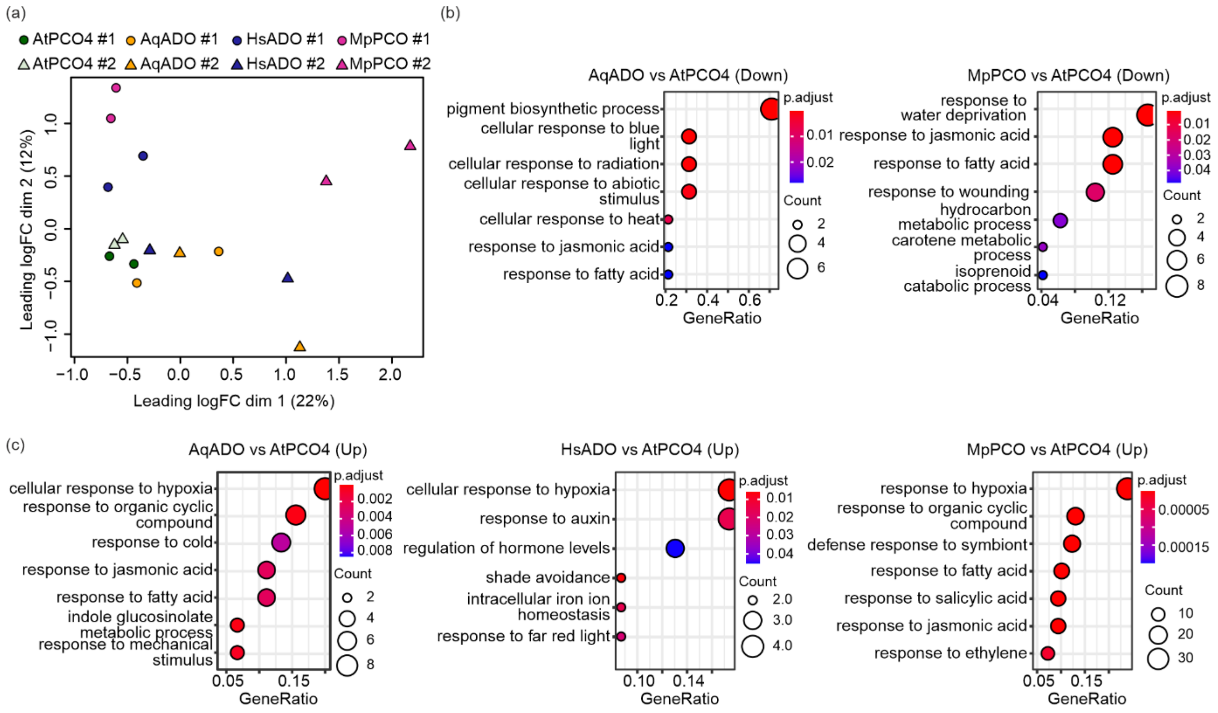

**Supplementary Figure 9 – Effect of NCO complementation on hypoxia transcriptional response and submergence tolerance.** (**a**) mRNA relative expression of three hypoxia-responsive genes (*ADH1*, *LBD41*, *PDC1*) in 24 hours of air (21% $O_2$ v/v), hypoxia (1% $O_2$ v/v) and 1 hour of reoxygenation (21% $O_2$ v/v). Statistical differences were analysed per condition using one-way ANOVA among treatments, followed by Tukey HSD post-hoc test ($n$ = 3, $p$ < 0.05). (**b**) Differences in survival rate between NCO-complemented plants after 2.5, 3 or 3.5 days of submergence or after 3.5 days at dark. Plants were categorized as 'alive', when presenting all green healthy leaves, 'damaged' when presenting at least one non-green leaf and 'dead' when bleached or fully wilted.

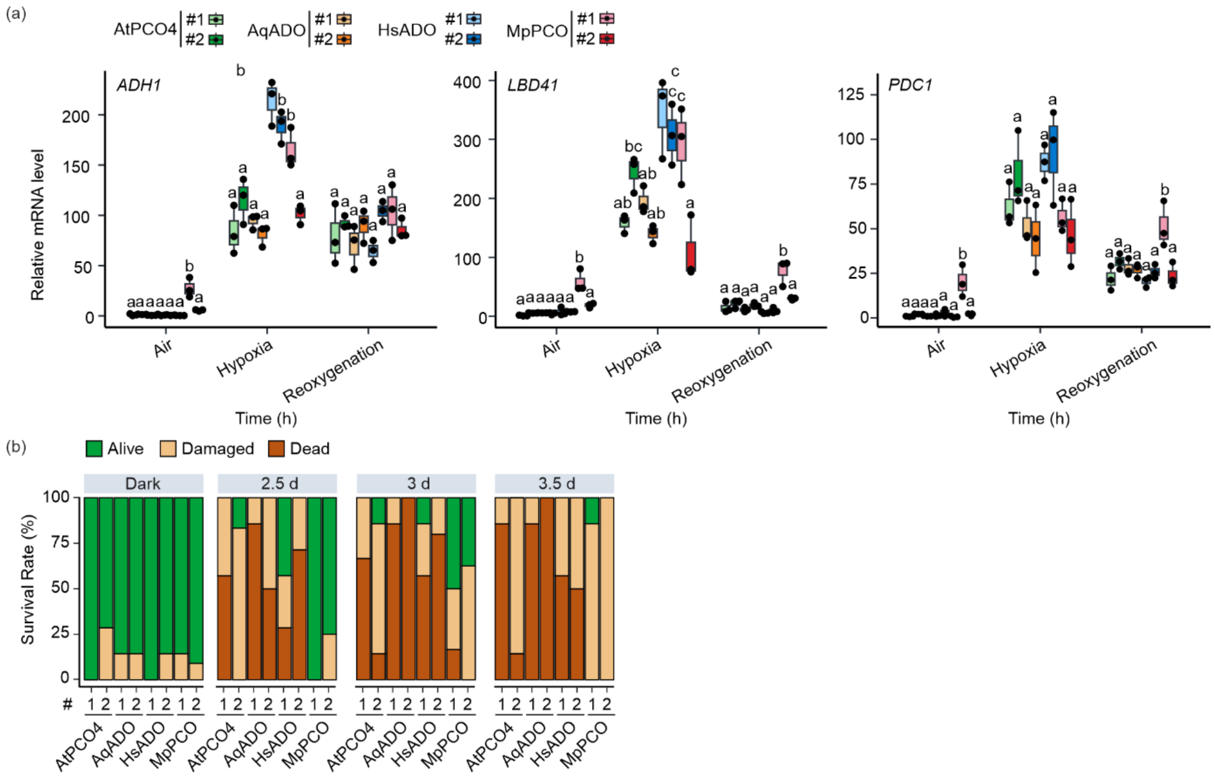

**Supplementary Table 1 - Steady-state kinetic parameters derived for NCO and activity toward $RAP2_{2-15}$ at atmospheric $O_2$.**

| | AqADO | HsADO | AtPCO4 | MpPCO | KnPCO |
|---|---|---|---|---|---|
| $K_M$ $RAP2.12_{2-15}$ (μM) | 205.7 ± 53.56 | 392 ± 91.61 | 690 ± 110 | 11 ± 1.1 | 914 ± 144 |
| $k_{cat}$ $RAP2.12_{2-15}$ ($s^{-1}$) | 15.51 ± 1.79 | 14.91 ± 1.99 | 31 ± 3.49 | 8.2 ± 0.5 | "18.0 ± 1.2 |
| $V_{max}$ $RAP2.12_{2-15}$ (μmol $min^{-1}$ $mg^{-1}$) | 36.66 ± 4.23 | 7.39 ± 0.99 | 63.4 ± 7.13 | 17.6 ± 1.1 | 36.6. ± 2.4 |
| $k_{cat}$/ $K_M$ $RAP2.12_{2-15}$ ($s^{-1}$/μM) | 0.08 ± 0.02 | 0.04 ± 0.01 | 0.04 ± 0.01 | 0.75 ± 0.09 | 0.02 ± 0.01 |

**Supplementary Table 2 - Steady-state kinetic parameters derived for NCO and activity toward $O_2$.**

| | AqADO | HsADO | AtPCO4 | MpPCO | KnPCO |
|---|---|---|---|---|---|
| $K_M$ $O_2$ (%) | 34.84 ± 3.86 | 81.5 ± 14.39 | 17.3 ± 1.69 | 3.2 ± 0.6 | 28.9 ± 4.1 |
| $k_{cat}$ $O_2$ ($s^{-1}$) | 22.67 ± 1.14 | 14.48 ± 1.55 | 23.8 ± 0.9 | 15.5 ± 0.7 | 59.3 ± 3.7 |
| $V_{max}$ $O_2$ (µmol $min^{-1}$ $mg^{-1}$) | 53.6 ± 2.69 | 29.2 ± 3.12 | 48.6 ± 1.83 | 31.2 ± 1.4 | 120.2 ± 7.6 |
| $k_{cat}$/ $K_M$ $O_2$ ($s^{-1}$/µM) | 0.05 ± 0.01 | 0.01 ± 0.01 | 0.11 ± 0.01 | 0.39 ± 0.08 | 0.17 ± 0.03 |

**Supplementary Table 3 – Reactions and reaction rates of mathematical model.**

| | Reactions | Reaction Rates | Description |
|---|---|---|---|
| $v_1$ | → RAP2.12 | $k_1$ | RAP2.12 synthesis |
| $v_2$ | RAP2.12 → null | $k_2 \cdot [\text{RAP2.12}]$ | RAP2.12 degradation |
| $v_3$ | RAP2.12 + $O_2$ + AtPCO4 → AtPCO4 | $k_3 \cdot [\text{AtPCO4}] \cdot \frac{[O_2]}{K_{M1}+[O_2]} \cdot \frac{[\text{RAP2.12}]}{K_{M2}+[\text{RAP2.12}]}$ | RAP2.12 degradation through oxidation by AtPCO4 |
| $v_4$ | RAP2.12 → mRNA + RAP2.12 | $k_4 \cdot \frac{[\text{RAP2.12}]}{K_{M3}+[\text{RAP2.12}]}$ | HRG mRNA production by RAP2.12 |
| $v_5$ | mRNA → null | $k_5 \cdot [\text{mRNA}]$ | HRG mRNA decay |
| $v_6$ | RAP2.12 + $O_2$ + AqADO → AqADO | $k_6 \cdot [\text{AqADO}] \cdot \frac{[O_2]}{K_{M4}+[O_2]} \cdot \frac{[\text{RAP2.12}]}{K_{M5}+[\text{RAP2.12}]}$ | RAP2.12 degradation through oxidation by AqADO |
| $v_7$ | RAP2.12 + $O_2$ + HsADO → HsADO | $k_9 \cdot [\text{HsADO}] \cdot \frac{[O_2]}{K_{M10}+[O_2]} \cdot \frac{[\text{RAP2.12}]}{K_{M11}+[\text{RAP2.12}]}$ | RAP2.12 degradation through oxidation by HsADO |
| $v_8$ | RAP2.12 + $O_2$ + KnPCO → KnPCO | $k_8 \cdot [\text{KnPCO}] \cdot \frac{[O_2]}{K_{M8}+[O_2]} \cdot \frac{[\text{RAP2.12}]}{K_{M9}+[\text{RAP2.12}]}$ | RAP2.12 degradation through oxidation by KnPCO |
| $v_9$ | RAP2.12 + $O_2$ + MpPCO → MpPCO | $k_7 \cdot [\text{MpPCO}] \cdot \frac{[O_2]}{K_{M6}+[O_2]} \cdot \frac{[\text{RAP2.12}]}{K_{M7}+[\text{RAP2.12}]}$ | RAP2.12 degradation through oxidation by MpPCO |

**Supplementary Table 4. Ordinary differential equations of the kinetic model. The reaction rates v1– v9 are given in Supplementary Table 3.**

| Model | Left-hand Sides | Right-hand Sides | Simulated Initial Concentrations (μM) |
|---|---|---|---|
| RAP2.12/AtPCO4 | d[RAP2.12]/dt | v1-v2-v3 | 0.00090 |
| | d[RNA]/dt | v4-v5 | 0.07230 |
| RAP2.12/AqADO | d[RAP2.12]/dt | v1-v2-v6 | 0.00990 |
| | d[RNA]/dt | v4-v5 | 0.08140 |
| RAP2.12/HsADO | d[RAP2.12]/dt | v1-v2-v7 | 0.00150 |
| | d[RNA]/dt | v4-v5 | 0.12210 |
| RAP2.12/KnPCO | d[RAP2.12]/dt | v1-v2-v8 | 0.00230 |
| | d[RNA]/dt | v4-v5 | 0.18710 |
| RAP2.12/MpPCO | d[RAP2.12]/dt | v1-v2-v9 | 0.00005 |
| | d[RNA]/dt | v4-v5 | 0.00390 |

**Supplementary Table 5. Parameter values used in the kinetic model.** Concentrations and the Michaelis-Menten constants (Kms) are given in μM, except Kms for $O_2$ are in %. Rate constants (ks) are expressed in $s^{-1}$, except $k_1$ is nM $s^{-1}$.

| Parameters | Description | Values | References |
|---|---|---|---|
| $k_1$ | RAP2.12 protein synthesis rate | 0.005 | (93, 94) |
| $k_2$ | RAP2.12 degradation rate | 0.0002 | (93) |
| $k_3$ | Catalytic rate constant for AtPCO4-mediated oxidation of RAP2.12 | 26.5878 | (40) |
| $K_{M1}$ | Michaelis-Menten constant for $O_2$ as a substrate of AtPCO4 | 16.4217 | (40) |
| $K_{M2}$ | Michaelis-Menten constant for RAP2.12 as a substrate of AtPCO4 | 270.2505 | (40) |
| NCOs | Protein concentration of AtPCO4, AqADO, HsADO, KnPCO and MpPCO | 0.1 | (95) |
| $k_4$ | Activation rate constant for HRG expression by RAP2.12 | 0.0016 | (95) |
| $K_{M3}$ | Michaelis-Menten constant for HRG expression | 0.05 | (40) |
| $k_5$ | HRG mRNA decay rate | 0.00038 | (96) |
| $k_6$ | Catalytic rate constant for AqADO -mediated oxidation of RAP2.12 | 153.4026 | Fitted |
| $K_{M4}$ | Michaelis-Menten constant for $O_2$ as a substrate of AqADO | 262.7200 | Fitted |
| $K_{M5}$ | Michaelis-Menten constant for RAP2.12 as a substrate of AqADO | 233.0448 | Fitted |
| $k_7$ | Catalytic rate constant for MpPCO -mediated oxidation of RAP2.12 | 12.1853 | Re-fitted (45) |
| $K_{M6}$ | Michaelis-Menten constant for $O_2$ as a substrate of MpPCO | 0.9715 | Re-fitted (45) |
| $K_{M7}$ | Michaelis-Menten constant for RAP2.12 as a substrate of MpPCO | 10.8438 | Re-fitted (45) |
| $k_8$ | Catalytic rate constant for KnPCO -mediated oxidation of RAP2.12 | 185.3389 | Re-fitted (45) |
| $K_{M8}$ | Michaelis-Menten constant for $O_2$ as a substrate of KnPCO | 63.5650 | Re-fitted (45) |
| $K_{M9}$ | Michaelis-Menten constant for RAP2.12 as a substrate of KnPCO | 2360.5 | Re-fitted (45) |
| $K_9$ | Catalytic rate constant for HsADO -mediated oxidation of RAP2.12 | 33.9905 | Fitted |
| $K_{M10}$ | Michaelis-Menten constant for $O_2$ as a substrate of HsADO | 123.8549 | Fitted |
| $K_{M11}$ | Michaelis-Menten constant for RAP2.12 as a substrate of HsADO | 156.6104 | Fitted |

**Supplementary Table 6 – List of primers for screening of transgenic lines.**

| Primer Name | Sequence |
|---|---|
| attB1 | GGGGACAAGTTTGTACAAAAAAGCAGGCTCC |
| attB2 | CACCCAGCTTTCTTGTACAAAGTGGTCCCC |
| pPCO4 Fw | ATGTGCGATTCATTTCCCAA |
| PCO4 Rv | GGGCTTTGAGCCGTCATCTCAGTATCCTTC |
| AqADO Rv | ATGATTTTCTTGAGCTTTTC |
| HsADO Rv | CGCCTCCGGGCCAGAAGCCG |
| SpADO Rv | GTGTTCGAAGACCTCACAGG |
| MpPCO Rv | GCCAGTCACAGCAGAATATGGA |
| KnPCO Rv | CGAGTCCAACATCAGCGCAA |
| LBb1 | GCGTGGACCGCTTGCTGCAACT |
| PCO5 Fw | GCCCATTTAGGTAGCTGCAGTG |
| PCO5 Rv | AGCTTCCTGTTCGAGACCAA |

**Supplementary Table 7 – List of primers used for TAIL PCR.**

| Primer Name | Primer |
|---|---|
| LAD1-1 | ACGATGGACTCCAGAGCGGCCGCVNVNNNGGAA |
| LAD1-2 | ACGATGGACTCCAGAGCGGCCGCBNBNNNGGTT |
| LAD1-3 | ACGATGGACTCCAGAGCGGCCGCVVNVNNNCCAA |
| LAD1-4 | ACGATGGACTCCAGAGCGGCCGCBDNBNNNCGGT |
| tail_RB0 | GGTGAAAAGGGCGAATTCTGCA |
| tail_RB1 | GGCGAATTCTGCAGATAAACTATCAGTGTTTG |
| tail_LB0 | GTAATGCATGACGTTATTTATGAGATGGGTT |
| tail_LB1 | TATCGCGCGCGGTGTCATCTATG |
| tail_AC1-A | AACACATCACGATGGACTCCAGAG |
| tail_AC1-B | AGAGTGCGACGATGGACTCCAGAG |
| tail_AC1-C | CCGTATATACGATGGACTCCAGAG |
| tail_AC1-D | CTTGGTTGACGATGGACTCCAGAG |
| tail_AC1-E | GAGATAACACGATGGACTCCAGAG |
| tail_AC1-F | GTACAGGAACGATGGACTCCAGAG |
| tail_AC1-G | TCTCGCCTACGATGGACTCCAGAG |
| tail_AC1-H | TGCTCCGAACGATGGACTCCAGAG |

**Supplementary Table 8 – List of primers used for qRT-PCR.**

| Gene | AGI Code | Forward Primer | Reverse Primer |
|---|---|---|---|
| *UBQ10* | *AT4G05320* | GGCCTTGTATAATCCCTGATGAATAAG | AAAGAGATAACAGGAACGGAAACATAGT |
| *LBD41* | *AT3G02550* | TGAAGCGCAAGCTAACGCA | ATCCCAGGACGAAGGTGATTG |
| *ADH1* | *AT1G77120* | TATTCGATGCAAAGCTGCTGTG | CGAACTTCGTGTTTCTGCGGT |
| *PDC1* | *AT4G33070* | CACAGAATCTTCAATGTTCTTACC | CCATGATAAAGCGTACATGGAA |
| *PCO4* | *AT2G42670* | CGAACCAGAGGATCCATCACAAGA | GCCGTCATCTCAGTATCCTTCACC |
| *AqADO* | *NA* | AGCCAGATCACCCCGAAGGA | AAGCGGCAGGAGCATCAGTC |
| *HsADO* | *NA* | CGGTGGTGGACAAAGGCCTA | AGCAGGTCCTTCAACAGCGT |
| *MpPCO* | *NA* | CACACACCAATGGTTGCCAG | GGGGTTGACCCAGTCATACG |
| *KnPCO* | *NA* | GACCTCAGCACCCTCCTAGA | AGGAGCAACAACATCGAGCA |

**Supplementary Table 9 – Multiple sequence alignment of NCO sequences in the eukaryotic kingdom using CLUSTAL Omega.**

```
Chytriomyces            -------------------------------------MVMIATALLAQRLHDILYGNTSG      23
Batrachochytrium        -------------------------------------------MPLLQAIGLFSERLFAQ      17
Rhizoclosmatium         --------------------------------------------MLLTAIGLVAHELFTN      16
Blyttiomyces            -------------------------------------------MSLLHTIGLLAHRLFSR      17
Spizellomyces           ---------MSPPCINESDSSPTTCCFPFCTEMENRRTQSLMDMSHLQRIGVLAYQVFAA      51
Chlamydomonas           ---------------------MQQNGTFASYNQTSSSGGSSGSSSTLQAFFSSAKAAVLR      39
Volvox                  ---------------------MHSQSAVAGS-QT--ANSQQTSGSRLQDFFTAARRAVLH      36
Ostreobium              -----------------------------AEEQGHGVGCETRTPLTAQRERRPCSAPAVR      31
Chlorella               -------------------------------------MPHRSPVRRLQFLMDVAQATFNQ      23
Picea                   -------------------------------------------MTAVQRLYDICKASFTS      17
Cryptomeria             ---------------------------------M-LEEQWFCTMTAVQRLYEICKASFTS      26
Taxus                   ----------------------------NIVFQH-CLELWFCTMTTVQRLYDSCKASFTS      31
Triticum                -------------------------------------------MAAIQKLYEVCKASLSE      17
Oryza                   ----------MPIASRRPLNGSSRRGWGWICLFV-FSRTSKCKMPKIKSLSNACKVSFSP      49
Zea                     -------------------------------------------MPKIKSLSNACRVSFSP      17
Arabidopsis             ------------------------------------------MPYFAQRLYNTCKASFSS      18
Camelina                ------------------------------------------MPYFAQRLYNTCKSSFSS      18
Nicotiana               -------------------------------------------MPAVQKLYNACKASLSP      17
Solanum                 -------------------------------------------MPAVQKLYNACKASLSP      17
Cannabis                ------------------------------------------MPYYIQRLFNTCKASFSP      18
Glycine                 ------------------------------------------MPYYIQRLYRLCNASFSP      18
Medicago                ------------------------------------------MPYCVQRLYRLCKASFSP      18
Klebsormidium           ------------------------------------------MASKVQKLYDACRAAFGA      18
Adiantum                --------------------MCLEEERLWPEPAE--DGSEQDPRPPIQRLYEFMTVFFEM      38
Selaginella             ----------------------------------------MTAMTAVQKLYEVCKASFSC      20
Ceratopteris            ----------------------MREESQSAPAVDESSVMMPEHAMAVQKLYESCRATFTR      38
Ceratodon               --------------------------------------MTMSSAPAIQRLYDVCKKVFTS      22
Physcomitrium           --------------------------------------MAMTSTPAVQKLYDVCKKVFTP      22
Diphasiastrum           -------------------------------------------MSAIQKLHDVCKASFSL      17
Marchantia              ------------------------------------------MMTAVQRLHDVCKATFTM      18
Sphagnum                --------------------------------------MGAMSTTAVQKLYDVCKATFSA      22
Caenorhabditis          --------------------------------------MDSPARHLQMAIRKIALLSRSF      22
Anopheles               ------------------------------------------MSALFARVFRQARQTFEY      18
Drosophila              ------------------------------------------MTTHFSNVLRQAFKTFDR      18
Candidula               ---------------------------------------MAAPILKLAQLSTKIFGVLSP      21
Octopus                 ------------------------------------------MALLIQEVSKAARRAFSG      18
Fasciola                ---------------------------------------MTTKIATVANLALHAFQQHSV      21
Schistosoma             ---------------------------------------MTTKIATVANLAFHAFQQHSA      21
Hydra                   ------------------------------------------MASLIQRIVRQAATTFPN      18
Daphnia                 ------------------------------------------MGSRIENVIRTAVQTFAT      18
Amphimedon              ------------------------------------MGVVK-MSGLVHQLVHQARRTFSR      23
Lamellibrachia          --------------------------------------KCSMNGTKIQRLAAQAFQTFSK      22
Strongylocentrotus      -------------------------------------------MAALQKVTRQAWDIFRL      17
Asterias                ----------------------------------------MANLQPIQKVAKLALETFKY      20
Scyliorhinus            ------------------------------------MPGDKMASSLIQTIARQARTTFKL      24
Xenopus                 ------------------------------------MPRDD-MSSLIQKVARQARQTFRS      23
Carassius               -----------------------------------MMPRDS-MTSTVQKIARQALTTFRN      24
Danio                   ------------------------------------------MTSTVQKIAKQALATFRN      18
Chelonia                MQGAVRAGAEGRALPRSTAESRGRAGSAERDYLGGTMPRDN-MASLIQRVAKQARITFRG      59
Homo                    ------------------------------------MPRDN-MASLIQRIARQACLTFRG      23
Mus                     ------------------------------------MPRDN-MASLIQRIARQACLTFRG      23
```

```
Chytriomyces         ADEGS--------------------------------------------------P-AGK   32
Batrachochytrium     PTPTH--------------------------------------------------PPPSA   27
Rhizoclosmatium      PSERA-------------------------------------------------------   21
Blyttiomyces         FPEPH----------------------------------------------------PQR   25
Spizellomyces        FPGRH----------------------------------------------------PEA   59
Chlamydomonas        GGLNE----------------------------------------------------TNT   47
Volvox               GTVNE----------------------------------------------------SNP   44
Ostreobium           TPATGTDAFP------------------------------------AGRLRNGLAGGPPC   55
Chlorella            PGGP------------------------------------------------------TR   29
Picea                SGPR------------------------------------------------------SP   23
Cryptomeria          NGPR------------------------------------------------------SP   32
Taxus                SGPR------------------------------------------------------SL   37
Triticum             KGPS------------------------------------------------------SP   23
Oryza                DGPI------------------------------------------------------SD   55
Zea                  EGPI------------------------------------------------------SE   23
Arabidopsis          DGPI------------------------------------------------------TE   24
Camelina             DGPI------------------------------------------------------SD   24
Nicotiana            NGPV------------------------------------------------------SE   23
Solanum              NGPV------------------------------------------------------SE   23
Cannabis             NGPV------------------------------------------------------SE   24
Glycine              NGPA------------------------------------------------------SE   24
Medicago             DGPV------------------------------------------------------SE   24
Klebsormidium        KGPA------------------------------------------------------SK   24
Adiantum             LIVVGGKG----------------------------------------------IDMFEQ   52
Selaginella          VQGVS-----------------------------------------------------AP   27
Ceratopteris         NAGSF-----------------------------------------------------PA   45
Ceratodon            T-SVP-----------------------------------------------------SE   28
Physcomitrium        A-SVP-----------------------------------------------------LE   28
Diphasiastrum        LGAVP-----------------------------------------------------SP   24
Marchantia           S-APP-----------------------------------------------------SA   24
Sphagnum             SSGVP-----------------------------------------------------SA   29
Caenorhabditis       D--------------------------------------------------------GSA   26
Anopheles            TTVS---------------------------------------------------DPKFA   27
Drosophila           AN-----------------------------------------------------HASFN   25
Candidula            E-------------------------------------------------------KITK   26
Octopus              ----------------------------------------------------SLKEPLFQ   26
Fasciola             HVTLASTAALTLGTSAGSTPGSGSLSNAQTSSHSPTDGSGAVPDSPNHAPRSASCPNGLP   81
Schistosoma          RA--------------ALSPASANINNGMGDSHPCTNGNSPTHSDKPVSPNLPVKHKQLS   67
Hydra                SL-------------------------------------------------------TVS   23
Daphnia              KPNGS-----------------------------------F-------------VNANVS   30
Amphimedon           EK-------------------------------------------------------TLP   28
Lamellibrachia       YR----------------------------------------------------RTDAFS   30
Strongylocentrotus   FFKD---------------------------------------------------RNANI   26
Asterias             VNKG--------------------------------------------------DKHVFL   30
Scyliorhinus         LPLA--------------------------------------------------ETGRFQ   34
Xenopus              GG-----------------------------------------------------GGPSA   30
Carassius            PSAVG-----------------------------------E-------------HGKVFL   36
Danio                PSVIG-----------------------------------E-------------HNKVFL   30
Chelonia             PA----------------------------------------------------PGPGFG   67
Homo                 SGGGR-----------------------------------GASDRDAASGPEAPMQPGFP   48
Mus                  SSTGS-----------------------------------E------------GPAPGFP   36


Chytriomyces         GRGDGATAMATPTPD-TDSLRVANPANTDTVANVLRLLLLKIALP---------------   76
Batrachochytrium     HTTARLLQSLTKLL---FKLKLESLA----------------------------------   50
Rhizoclosmatium      ADLEEALTLLLERLT-PAMLKL--------------------------------------   42
Blyttiomyces         EVIAEKLGCLVKLVT-TEQLNLQHVK----------------------------------   50
Spizellomyces        DAILDRLCSLIRDVT-LGHLKVKP------------------------------------   82
```

```
Chlamydomonas           ETVNSLL-RLMSHIS-LEELGLDPDMVQDPFLGALKLH----------------------      83
Volvox                  DTVAQLA-NMMGAIR-LEELGLDSDMVADPFLGALKLH----------------------      80
Ostreobium              VAKDQAPAIPPEPAM-DDGQERRPRLQELYDVGWRTFGRRGGGAPSREEVLAVLGALREV      114
Chlorella               QQVEALQ-RTLSLVP-LEELGLSPQQREQQQHEVHSSSSSRSCSPGLGDGAHMLHDELER      87
Picea                   DVLQNVR-DILDTIK-PSDVGLEEEALV-RGWSGPAFG----------------------      58
Cryptomeria             EILQNVH-DVLDTIK-PSDVGLEEAALVAHRWSGSSHG----------------------      68
Taxus                   EFLQNVR-DVLDTIK-ASDVGLEEGELVARGWSGSPYG----------------------      73
Triticum                EAVENVR-AVLDMIT-PSDVGLECEAQAVRVWRRPRVL----------------------      59
Oryza                   EALERVR-ALLDEIR-PIDVGLDNEAQIARNWNNSTRQ----------------------      91
Zea                     EALERVR-ALLDMIR-PLDVGLDNEAQIARNWSSSTRP----------------------      59
Arabidopsis             DALEKVR-NVLEKIK-PSDVGIEQDAQLARSRSGPLN-----------------------      59
Camelina                EALEKVR-NVLEKIK-PSDVGIEQEAQLARTRSGPLN-----------------------      59
Nicotiana               DALEKVR-SLLDKIK-PSDVGLEQEAQLVRSWNGTLH-----------------------      58
Solanum                 DALEKVR-SLLDKIK-PSDVGLEQEAQLVRSWNSTLH-----------------------      58
Cannabis                ESLEKVR-AMLEKIK-PSDVGLEQEAQLVRNWPGTLP-----------------------      59
Glycine                 EAIEKVR-EKLERIK-PSDVGLEQEAQVVRNWSGSML-----------------------      59
Medicago                EVVKKVR-EKLDRIK-PSDVGLEQEAQVVRNMSRTVL-----------------------      59
Klebsormidium           ENLQLLQ-EALDSVT-CADVGLDPVTDG-ETRGFGFFG----------------------      59
Adiantum                RHRECKESLMADNIK-PIDVGLDESMPPHNDRGFGFFG---------------------M      90
Selaginella             EAVARVR-AVLDDIK-CSDLGLQEDNCQRNGRST--------------------------      59
Ceratopteris            QGLQRVR-SILDSIK-PSDVGLDENAPIQQNSRTNLLG---------------------R      82
Ceratodon               AQVSNVR-AVLDTIK-PQDLGLTIQSDQQNR---GDRG---------------------F      62
Physcomitrium           AQVAKVR-AVLDSIK-PQDLGLGTIDQHN-------RG---------------------L      58
Diphasiastrum           PAIQTVR-NALDNIE-CADVGLDWKEAQQDDDGFGYYG---------------------L      61
Marchantia              QALQRVR-TILDSIQ-ASDVGLDEVDAMSQERGFGFFG----------------------      60
Sphagnum                AALERVR-LALENVK-ALDVGLEEEEQP--ERGFGFFG----------------------      63
Caenorhabditis          ESINIIQ-SIVNNFDSRIFQRLIPGRISRVIQT---------------------------      58
Anopheles               TNLTALR-RLVDQLT-LADIGLDGSLVATDTFQ---------------------------      58
Drosophila              ANLQHLR-QLTDELT-YRDLHLREELFRNVG-----------------------------      54
Candidula               DHLSPLI-RAMNNIS-SEDVYFDPKEIEERDAEL--------------------------      58
Octopus                 EKLNELV-KLMNQVR-GRDLNFNPALTERSKNN---------------------------      57
Fasciola                VALQRLL-ISVHNLK-PKDIGFDTSWVSDSQVY---------------------------      112
Schistosoma             ESLSRLL-ACVRSLT-PKDVGFDIRWISDSNIS---------------------------      98
Hydra                   SNLENLI-ALVEGLT-ANDLGLKADAKDALA-----------------------------      52
Daphnia                 HNMNILR-QLLSEICPADDLGLTRYSGQPRFPL---------------------------      62
Amphimedon              EEMEKLK-KIMSQIT-PKDFNLDSTAIDKPWDY---------------------------      59
Lamellibrachia          EQLTNLR-DLMNHIT-KVEVNFNSQLVKNKEVY---------------------------      61
Strongylocentrotus      DGLKALQ-ETMSKIQ-ASDLNIDASRAMIEQQRLPPNPA---------------------      63
Asterias                DKLESLC-ASVNRIR-AADVGVDLTANALP------------------------------      58
Scyliorhinus            DNLQQLN-SLLEKIQ-VDDLNLGPRRNQG-------------------------------      61
Xenopus                 ESLGQLR-KLVSQVR-AEDVGLGVSRSS--------------------------------      56
Carassius               ENLSKLK-SLVAEVR-AADLRIAPRTSASAPV----------------------------      66
Danio                   ENQSKLK-SLLAEVR-AADLKIAARTPESAPV----------------------------      60
Chelonia                ENLHRLQ-QLLNEVQ-AEDLHLAPRGPAAVPG----------------------------      97
Homo                    ENLSKLK-SLLTQLR-AEDLNIAPRKATLQ------------------------------      76
Mus                     ENLSLLK-SLLTQVR-AEDLNIAPRKALPQ------------------------------      64


Chytriomyces            ---------MLSVQIPSVNSIEYYHIWGSE-SMTMCLFVLKKGSTLPIHSHPN------M      120
Batrachochytrium        -------------ILSPQHPIEFYPLYETK-DFAICIFVLKKGTTMPTHDHPG------M      90
Rhizoclosmatium         -------------EKPPSSPIEYSNIYDCP-AFNIAAFVIRKGETMPIHDHPN------M      82
Blyttiomyces            ----------------VIDPVEYYNIYESD-VFTLCVFALKGGTTMPTHDHPQ------M      87
Spizellomyces           ----------------AQYPVEYYPIYESS-VYTMCVFVLKRGTTMPTHDHPK------M      119
Chlamydomonas           ---------------LRDSRIKYMRIYEDHT-LTFGLFCFPAGATIPLHNHPG------M      121
Volvox                  ---------------LRDSRIKYMRIYEDPS-LTFGLFCFPAGTVIPLHNHPDGHSANPS      124
Ostreobium              PIEEVGLPGPLGRRGRSARSITYLPVKEDRR-LHLGVFCMPPRATIPLHDHPG------M      167
Chlorella               ERGRPYPCTESEA-LDPRPPITYLLIYEDAR-VSLGIFCLPARAKIPLHNHPG------M      139
Picea                   INGRRH---------RYMPPITYLHIHECER-FSIGIFCMPPSSIIPLHNHPG------M      102
```

```
Cryptomeria             VNGRRGRNGSP----RYFSPITYLHIHECEN-FSIGIFCMPVSSVIPLHNHPG-----M      117
Taxus                   INGRKGRNGGL----RYTSPITYLHIHECEK-FSIGIFCMPASSTIPLHNHPG-----M      122
Triticum                -----NRKTVL----HSSPAIRYRHIYECKS-FSIGIYCIPASSIIPLHNHPG-----M      103
Oryza                   PNGRRGRNGAN----QFTSPIKYLHIHESES-FSMGIFCMPPSSVIPLHNHPG-----M      140
Zea                     SNGRRGRNGAN----QFAAPIKYLHIHECES-FSMGIFCMPPSSVIPLHNHPG-----M      108
Arabidopsis             -----ERNGSN----QSPPAIKYLHLHECDS-FSIGIFCMPPSSMIPLHNHPG-----M      103
Camelina                -----ERNGAL----QSPPAIKYLHLHECDS-FSIGIFCMPPSSMIPLHNHPG-----M      103
Nicotiana               -----DRNGGI----RSPPPIKYLHIHECDS-FSMGIFCMPPSSIIPLHNHPG-----M      102
Solanum                 -----ERNGEL----QPPPPIKYLHLHECDS-FSMGIFCMPPSSIIPLHNHPG-----M      102
Cannabis                -----ERNGRH----QSLPPIKYLHLHEDDS-FSIGIFCMPPSSIIPLHNHPG-----M      103
Glycine                 -----EHNGSH----QSLPPIKYLHLHECDS-FSIGIFCMPPSSIIPLHNHPG-----M      103
Medicago                -----EQNGSH----HSLPAIKYLHLHECDS-FSIGIFCMPPSSIIPLHNHPS-----M      103
Klebsormidium           RQGPWGRHSAAP--PRWAPPISYQHIYEDQH-ISIGIFCLPTCASIPLHNHPG-----M      110
Adiantum                FNGQKGRHSSMV--SRWAPPISYKHIYECQE-FSMGIFCLPTSAVIPLHNHPG-----M      141
Selaginella             --------DQPP--ESAAPAITYLHLYECNK-FSMGIFCLPQSAALPLHNHPG-----M      102
Ceratopteris            INGRRGK---------VSPAISYLHLYECNY-FSMGIFCLPKSAVIPLHNHPG-----M      126
Ceratodon               VDDVQGNWRSIL--PRYAPPISYLHLYECDL-FSMGIFVLPTSASIPLHDHPG-----M      113
Physcomitrium           GDDGKEKCRPIL--SRYVPVISYLHLYECDL-FSMGVFVLPASASIPLHNHPG-----M      109
Diphasiastrum           VKGCRGRLSAMV--PRWAQPITYLHLYEKENFFSMGIFCLPTSAAIPLHNHPG-----M      113
Marchantia              TNGRRGRHTPMV--ARWAPPISYLHLYECED-FSMGIFCLPTSAQIPLHNHPG-----M      111
Sphagnum                SNGETGRHSSMV--PRWSPPISYLHLYECEQ-FSMGIFCLPTSAAIPLHNHPG-----M      114
Caenorhabditis          ----------L---RKETSGIRYADVYQDDYCHVNTFGLLKSGLKIPLHDHPN-----Q      99
Anopheles               --------------RPTKAPCTYVGVFEND-CFAMSVFVLRENYTMPLHDHPR-----M      97
Drosophila              ---------------SHRAPCSYMHIFEDD-RFSMSLFIVRGASTIPLHDHPM-----M      92
Candidula               ---------QK---LGEVAPVTYMHIHQDH-NFTMAVFVVKAGSVLPLHDHPK-----M      99
Octopus                 -----------------NAPVTYIHIFEDE-VYSMGIFVLKENSRIPLHDHPG-----M      93
Fasciola                -----------------AAPVAYVHITENE-VFSMGIFILRSGSRIPLHDHPG-----M      148
Schistosoma             -----------------SAPVAYVHIMENE-VFSMGIFILKPGSRIPLHDHPG-----M      134
Hydra                   --------------IYPQAPVTHVSIYEGK-NFTMGVFILHPGMAIPLHDHPG-----M      91
Daphnia                 --------NLF---SSKAAPVSYMEIFENQ-TVSIGIFVLKDGASIPLHDHVG-----M      104
Amphimedon              ---------PP---FVTDAPAAFMNVYECS-EFNVAIFMLKANKEMPLHDHPE-----M      100
Lamellibrachia          --------QAE---SGGKPPVTYVHLWEDD-VFSMGIFVLRNGAKIPLHDHPG-----M      103
Strongylocentrotus      --AFEAAAGGD---PSQRAPVGYMHIFEDG-VMTMGVFIIREGSRIPLHNHPG-----M      111
Asterias                ----------P---VPATAPVGYMFIAENS-AVSMGVFVVERGCRLPLHDHNG-----M      98
Scyliorhinus            --------------PRHGPPVTYMHICETE-YFSMGVFLLQGGACIPLHDHPG-----M      100
Xenopus                 --------------GGKGPPVTYMHICETS-CFSMGVFLLRPGACIPLHDHPG-----M      95
Carassius               ---------PS---QRVAPPVTYMHIYETD-TFSMGVFLLKTGASIPLHDHPG-----M      107
Danio                   ---------PH---QRIAPPVTYMHICETD-SFSMGVFLLKTGASIPLHDHPG-----M      101
Chelonia                --AGCSRAGAG---AGAVPPVSYMHICETE-SFSMGVFLLRSGACIPLHDHPG-----M      145
Homo                    ---------PL---PPNLPPVTYMHIYETD-GFSLGVFLLKSGTSIPLHDHPG-----M      117
Mus                     ---------PL---PRNLPPVTYMHIYETE-GFSLGVFLLKSGTCIPLHDHPG-----M      105
                                             .  :            .     :* *.*

Chytriomyces            TVFSRVIHGDLHVKTYHLEGDSHANTSSSNSRSSLPTASASEGVTPASSPRLFQTTTQPS     180
Batrachochytrium        TVFTKLISGDMHVKTFELINDSNSILKNAKCQL---------------------------     123
Rhizoclosmatium         TVFTKLIHGDLHVKTFELMDETGPVRRARVITDKICTASTTTPSNSFTSSSNQHSSPSAH     142
Blyttiomyces            MVFSKVIRGDLRVRTYQFAEDESGPHT-VV-DPESGELVKVRSAHAS-------------     132
Spizellomyces           TVFSKIVSGDLHVRTYEFAKGSTAQAHAVV-DPRTGDLVQARPTKAG-------------     165
Chlamydomonas           TVFSRLLFGRLRVSAFDWVVQPPEPIT--EG-----------------------------     150
Volvox                  RSATRLLFGQLRVSAYDWVVQPAAPIT--SG-----------------------------     153
Ostreobium              TVLSRVLYGRVRITSYDWAEGAGGGAG---------------------------------     194
Chlorella               TVLSRVLYGDMHVCSYDWAEPGDADA---VE-----------------------------     167
Picea                   TVLSKLLYGSMHAKAFDWINPSEAAD---TS-----------------------------     130
Cryptomeria             TVLSKLLYGSVHAKAYDWIDPSEVSD---SS-----------------------------     145
Taxus                   TVLSKLLYGSMHAKAYDWNDPSEVAD---SS-----------------------------     150
Triticum                TVLSKLLYGKVHVKAYDWIDIEEPGN---LS-----------------------------     131
Oryza                   TVLSKLLYGTLHAESYDWIDVTDPTDQLQEL-----------------------------     171
Zea                     TVLSKLLYGRLHAESYDWVDIPDHPI--DQL-----------------------------     137
```

```
Arabidopsis              TVLSKLVYGSMHVKSYDWLEPQLTEPE-DPS----------------------------     133
Camelina                 TVLSKLVYGSMHVKSYDWLEPELTEPD-DQL----------------------------     133
Nicotiana                TVLSKLVYGSFHVKAYDWIDIPGPSV--LTE----------------------------     131
Solanum                  TVLSKLVYGSFHVKAYDWINIPGPLV--PLE----------------------------     131
Cannabis                 TVLSKLIYGSLYVNSYDWLDLPRPDD--P-F----------------------------     131
Glycine                  TVLSKLLYGSMYVKSYDWIDAPGSND--P-S----------------------------     131
Medicago                 TVLSKLIYGSLYVRSYDWIDVPGFTY--S-S----------------------------     131
Klebsormidium            VVMSKLLYGDMHVRSYDWANQEAPPVGPQHP----------------------------     141
Adiantum                 TVFSKLLYGSMHVKAYDWVDLN------KVL----------------------------     166
Selaginella              TVLSKLIYGSMHVKSYDWVDPSDHTA---SS----------------------------     130
Ceratopteris             TVMSKLLYGSMHVKAYDWVDHN------DVS----------------------------     151
Ceratodon                TVLSRLLYGNMHVRAYDWVEPHDGRLNRNPA----------------------------     144
Physcomitrium            TVLSRLLYGSMHVRAYDWVDPHDEKLNRNPS----------------------------     140
Diphasiastrum            TVFSRLLYGSMHVRSYDWVNPFSELSG-DAS----------------------------     143
Marchantia               TVLSKLLYGSMHVKAYDWVNPCDEKLNSDPS----------------------------     142
Sphagnum                 TVLSRLLYGSVHVRAYDWVNPFDEKLNADPS----------------------------     145
Caenorhabditis           HAVMKVYQGSVKIRSFSIIDCDESDDDDEGGGAASSSSA---SPVRV------------     143
Anopheles                HGLLRVVSGAVQICSYSELARRDADELLVGV--------------GK------------     130
Drosophila               FGLLRCIWGQLMVDSFSHQLGPDEPLTYDP-----------------------------     122
Candidula                HGLLKVLYGKVRISSYTEVDQKPLPENIARLQVSQS-------QKRS------------     139
Octopus                  FGIIKVIHGTLSLESFDEVPVSSEQPTSSASTTATSFNN---EFKKF------------     137
Fasciola                 YGILRVMYGSLRCRSFTPLQNLKPSHPAYPRSSLTEAFP---GSG-W------------     191
Schistosoma              YGILKVLTGSVRCRSFTRLKNVKSTTDLGNYKPSLSVFD---DSK-W------------     177
Hydra                    NGICKVLYGSIKLTSFEGLQSRNFMKGGTS-----------------------------     121
Daphnia                  YGILKVLYGSLNVQSYSSIDLPGQTNSSIVH----------------------------     135
Amphimedon               HGLMKILSGSMCVTSYNRPEKRGSNSY--------------------------------     127
Lamellibrachia           HGLIKVLEGKAKLNYYSDLSVNEIPVE---------VLP---QLHPW------------     138
Strongylocentrotus       HGLLKVLYGDISVRTFNTITEDWTKFPISKF-EGFPEN------EPP------------     151
Asterias                 HGVIKVLSGTIRMHSYDLLTEDEVNSIKIPD-SLAHPNS---ESDEE------------     141
Scyliorhinus             HGLLKVLYGSVAVKCFDKEETEPGAGP---A-AGVQFNP---PLLQC------------     140
Xenopus                  HGLLKVLYGKLRISGFDRLEASEPP-------EALAFSP---PLLPY------------     132
Carassius                HGMLKVIYGKVRISCFDRLDKPRD-----GA-SGAHFSP---PLVPF------------     145
Danio                    YGMLKVIYGKVRISCFDRLDKPRD-----GA-SGVQFNP---PLMPF------------     139
Chelonia                 NGMLKVLYGSLRIACLDPLPPAAPPGA--------------------------------     172
Homo                     HGMLKVLYGTVRISCMDKLDAGGGQRPRALP-PEQQFEP---PLQPR------------     160
Mus                      HGMLKVLYGTVRISCMDKLDTGAGHRR--PP-PEQQFEP---PLQPL------------     146
                             :   *

Chytriomyces             SHAQHPNLPHNARLHRDE---IISH------TSPGLTSILEINSNTPNLHTFTAVS-DHV     230
Batrachochytrium         ------------------------------------------------------------     123
Rhizoclosmatium          PPSPALTH--------------ASE------PYSTETIFKIFPSDGPNLHSYSAVS-DYA     181
Blyttiomyces             -----RTR--------------TIP------STSPDSLLVIRPDGGPNMHSFTAAS-PEV     166
Spizellomyces            -----LNR--------------IIS------ASDPESLLLIRPEGGPNMHSFTAVS-EEV     199
Chlamydomonas            -------SGHRAQLVYDA---TVEAS---------SPPLVLFPASGGNLHEFTAET--PC     189
Volvox                   -------SA-RARLVCDA---VIGPA---------TPPLVLFPSSGGNLHEFTAET--PC     191
Ostreobium               --------GGAARRVADE---VLSGP----------TGTSLLTPVHANVHSLEALT--AC     231
Chlorella                -------LPRAARRVVDS---TLTAA---------DQPTVLFPDAGGNIHEFTALT--DC     206
Picea                    -------KAKLAKLVKDC---EMSAP---------CDTTILYPTSGGNLHSFRALT--PC     169
Cryptomeria              -------KAKLAKLVKDC---EMSAP---------CDTTILYPTYGGNIHSFRALS--PC     184
Taxus                    -------KAKLAKLVKDC---EMSAP---------CDTTILYPTRGGNIHSFRAMS--PC     189
Triticum                 -------KVRPAKVVRDG---EISAP---------CAAMVLRPTGGGNVHALKAIT--PC     170
Oryza                    -------SVRPARLVRDR---EMSAP---------E-TTILYPNRGGNIHTFRAIT--PC     209
Zea                      -------QTRPAKLVKDC---EMTAP---------E-TTILYPNAGGNIHTFRAIT--PC     175
Arabidopsis              -------QARPAKLVKDT---EMTAQ---------SPVTTLYPKSGGNIHCFKAIT--HC     172
Camelina                 -------QARPAKLVKDK---EMTAP---------TPATTLYPKSGGNIHCFKAIT--HC     172
Nicotiana                -------GTRPAKLVKDC---DMTAP---------CGTTVLYPTSGGNIHCFKAIS--PC     170
Solanum                  -------GARPAKLVKDC---DMTAP---------CGTTALYPTSGGNIHCFKAIT--PC     170
Cannabis                 -------EARPAKLVRDT---EMIAP---------CPATILYPTSGGNIHCFRALT--PC     170
```

```
Glycine              -------EARPAKLVKDT---EMTAP---------SPTTVLYPTSGGNIHCFRAIT--PC   170
Medicago             -------GARPAKLVKDT---EMTAP---------SPTTILYPTNGGNIHCLRAIT--PC   170
Klebsormidium        -------PRRNAVLVCDE---VLSAP---------CDALVLYPTSGGNIHAFTGVT--PC   180
Adiantum             -------QPRLAKLVVDR---TMSAP---------CEPAVLYPTSGGNMHAFTAVT--SC   205
Selaginella          -------TPRLAMPVKDH---VLVAP---------CDTAVLYPTSGGNIHSFTAVT--SC   169
Ceratopteris         -------HPRLAKLVVDR---TLTAP---------CESSILYPKSGGNMHVFTAIT--PC   190
Ceratodon            -------IARQAKLVVDH---VLVTGPNETEPLRAPATEVLYPTSGGNIHSFTALT--SC   192
Physcomitrium        -------IARQAKLVIDH---VLGTEPNGTEPQQTSATEVLYPTSGGNIHAFTALT--SC   188
Diphasiastrum        -------TPRLAKLVADH---VLTAP---------CESAVLFPASGGNIHAFTALT--PC   182
Marchantia           -------VPRSAKLVVNR---VLTAP---------CETATLHPTSGGNIHAFTAVT--PC   181
Sphagnum             -------RPRLAKLVVDH---VLTAP---------CETAVLYPTTGGNIHAFTALT--PC   184
Caenorhabditis       ---RYE-GEIVLS--------SGSEQ----------QHSAVLGPRNGNIHEVTSLE--PH   179
Anopheles            ---GQR-HVLVAAEPEKI---ISSSP----------GDCALLTPTERNFHEITAIG-GPA   172
Drosophila           ----HQ-TVVKVNVEEPK---LVTPA----------SPCATLTPRKRNYHQIAQIGSGVA   164
Candidula            ---NSC-IIKTVYRHNDI---VLSET----------DECCVLSPSVGNFHEIQPET-KFA   181
Octopus              ---LRD-YDRTPKPFLEVPKKKASVN----------DDCCVLSPTKGNLHEVRPDG-GPA   182
Fasciola             ---QFR-DLIVARPHQDV---VLNVE----------SCAQLLTPTEGNLHEVTPVD-GPA   233
Schistosoma          ---QLT-DLIIARPHQDT---VLSPD----------HQPCLLFPIEGNLHEITPVD-GPA   219
Hydra                -------KYVQVKRIPEK---ILTAD----------TKSQFFLPIREIYHSMKATD-GPA   160
Daphnia              -----QPQYLKARRFPIS---CINEK----------DSPAVLSPSEHNLHTIWTVG-GPA   176
Amphimedon           ----------IARFFRSI---LATPA----------TEPCHFTPEEGNIHQIKAQD-SHV   163
Lamellibrachia       ---QIQ-MTRVVSKCKEE---EVNAD----------DVPCLLTPDEGNVHEVCAVD-GPM   180
Strongylocentrotus   ----KKHLLAPTRLGIDQ---HFTAS----------SEAVLLTPREGNYHSLESVG-GPA   193
Asterias             ---LMRNLIPVRKKAEAK---IITPE----------SKPCVLSPMESNVHSLESID-GPA   184
Scyliorhinus         ---QQA-TLRRSLQSSTK---VLDQA----------SGPCILTPLHNNMHQIDALD-GPA   182
Xenopus              ---QYN-CVCRAALRSAG---DFGDN----------SPPCLLAPHRENLHQISAEE-GPA   174
Carassius            ---QSS-SLRPAVLRSVG---EYTEE----------NGPCVLSPQKDNIHQIDAVD-GPT   187
Danio                ---QRG-SLRPSVLKSVG---EFTED----------SSPCVLSPQQDNIHQIDAVD-GPT   181
Chelonia             --------RRRALLRSRQ---LYTPA----------SPPCLLSPHTDNLHQIDAVD-GPA   210
Homo                 ---ERE-AVRPGVLRSRA---EYTEA----------SGPCILTPHRDNLHQIDAVE-GPA   202
Mus                  ---ERE-AVRPGVLRSRA---EYTEA----------SGPCVLTPHRDNLHQIDAVD-GPA   188


Chytriomyces         VVFDLIFPPYDE-------------------------DARMCSYYQERHDSANISG----   261
Batrachochytrium     ------------------------------------------------------------   123
Rhizoclosmatium      IIVDIMGPPYAV-------------------------GEREITYFKEVPLSNNNT--TEF   214
Blyttiomyces         MILDVIGPPYNL-------------------------TDRPCTYYRELARRAGTGGRQRR   201
Spizellomyces        VLLDIIGPPYNS-------------------------EDRPCTYFREVSWDEGC-----K   229
Chlamydomonas        AVLDLLTPPYEP-----------------------P--TRNCTYYRLQRPPMVWPVAR--   222
Volvox               AVLDLLTPPYEP-----------------------P--ERNCTYYRVQRQAASWPVGL--   224
Ostreobium           AVLDVLAPPYAP-----------------------S-KGRDCTYFRECPGGDA-------   260
Chlorella            AVLDLMSPPYST-----------------------E-DGRDCTYYSIVSEEQQ-------   235
Picea                ALLDVLAPPYST-----------------------D-NGRHCSYYRKLPKR-IPSG----   200
Cryptomeria          ALLDVLAPPYST-----------------------E-EGRHCTYYRKSFRK-VPSG----   215
Taxus                ALLDVLAPPYST-----------------------D-EGRHCTYYRKSLRK-VPPG----   220
Triticum             AILDILSPPYSS-----------------------K-DGRHCSYFRRRQKS-HPTG----   201
Oryza                ALFDVLSPPYSA-----------------------E-KGRDCSYFRKSSVRETPPV----   241
Zea                  ALFDVLSPPYSA-----------------------E-DGRHCSYFRKSQMN-QPPV----   206
Arabidopsis          AILDILAPPYSS-----------------------E-HDRHCTYFRKSRREDLPGE----   204
Camelina             AILDILSPPYSS-----------------------E-HDRHCTYFRKSRREDLPGE----   204
Nicotiana            AIFDILSPPYSS-----------------------E-DGRHCTYFRRSPIGDLPGE----   202
Solanum              AIFDILSPPYSS-----------------------E-DGRHCTYFRRSPVADLPGE----   202
Cannabis             AIFDILSPPYSS-----------------------E-NGRHCTYFRTSPRKDLPGS----   202
Glycine              AIFDILSPPYSS-----------------------D-HGRHCTYFRRSQRKDLPVN----   202
Medicago             AVFDILSPPYSS-----------------------E-DGRHCTYFRQSQRKDLPVN----   202
Klebsormidium        AVLDVVAPPYDP-----------------------E-SGRHCTYYRETPVPAMVGK----   212
Adiantum             AVLDILAPPYSA-----------------------Q-DGRHCTYYRDFPYSSFAS-----   236
Selaginella          AVLDVLAPPYNT-----------------------T-AGRPCVYYRLAAATSP-------   198
```

```
Ceratopteris            AVLDVLAPPYSA-----------------------E-DGRHCTYFRALPYASRTG-----    221
Ceratodon               AVLDVLAPPYSP-----------------------G-TGRHCTYYRATSNDGV-------    221
Physcomitrium           AVLDVLAPPYAP-----------------------A-TGRHCTYYRITSHEGE-------    217
Diphasiastrum           AVLDVLTPPYSP-----------------------I-DARHCIYYRESKFCGYPH-----    213
Marchantia              AVLDVLAPPYSA-----------------------V-TGRHCTYYREAHSVGSN------    211
Sphagnum                ALLDVLAPPYSP-----------------------A-TGRHCTYYHNIPRAVFSGSD---    217
Caenorhabditis          TYFCDFFIANSP----------------------------NCNYYRVDESSEPL------    205
Anopheles               AFFDILSPPYNTASQPQYYFYRKVPVPRHLAELEPI-GSPDKPSPLHD------------    219
Drosophila              AFFDILSPPYDAD-------------------MPTY-GPRQCRFYRAT------------    192
Candidula               AFLDVLAPPYEG-----------------------D-CDSGCHYYKDLNPG---------    208
Octopus                 AFLDILAPPYDT-----------------------D-SDRRCNYYEITKGHSPP------    212
Fasciola                VFLDILAPPYDH-----------------------DLGTRECRFYKEVILLGNHVPTRPV    270
Schistosoma             VFLDILAPPYDH-----------------------DLGTRECRFYKEVIIP-QMNSNALL    255
Hydra                   AFFDILAPPYRT----------------------KD-YKTDCHYFRELTVSEHPEIDLEK    197
Daphnia                 SFLDILAPPYDPSGT-----------------LNGG-EVRDCYYYSDVAGEDGMP-----    213
Amphimedon              AFLDILAPPYAP-----------------------E-EGRDCTYYFKDTKQSN-------    192
Lamellibrachia          AFLDILSPPYDN--------------------------VRKCHYYKTLSPVSTDNRTT--    212
Strongylocentrotus      AFLDILSPPYDP-----------------------V-IGRDCQYFKELKSLQPSS-----    224
Asterias                AFLDVLSPPYDP-----------------------D-TGRDCVYYTETPFSSSNLKL---    217
Scyliorhinus            AFLDILAPPYDP-----------------------E-XGRDCHYFKVLQTVSEGPGKA--    216
Xenopus                 AFLDILAPPYDP-----------------------A-DGRDCHYYQLIHPAAPPGETD--    208
Carassius               AFLDILSPPYDP-----------------------D-EGRDCHYYKVLHAHSEASDR---    220
Danio                   AFLDILAPPYDP-----------------------D-EGRDCHYYKVLQAHSEAADK---    214
Chelonia                AFLDILAPPYDP-----------------------E-NGRDCHYYRLLEAPAG--PA---    241
Homo                    AFLDILAPPYDP-----------------------D-DGRDCHYYRVLEPVRPKEAS---    235
Mus                     AFLDILAPPYDP-----------------------E-DGRDCHYYRVVEPIRPKEAS---    221
```

```
Chytriomyces            -----------DVVTTVATELSRMESSALIGKEQNAELHASISRLSVHEDGY-SSESRED    309
Batrachochytrium        ------------------------------------------------------------    123
Rhizoclosmatium         VVENGSTPSSAALFDSFSSSSSDSCIPTTGSVKRKRSETGLSDDAADQVVAK-KQTSS--    271
Blyttiomyces            RASGGVVRAGPPQVAGLASVSALGEAAPVASREDTAAGSAPATVPTRRAHPI-PPVSAAA    260
Spizellomyces           LVAMGVEATAPG----KKSKNKKKKKRRRAVGPERKTDAMPIQENGSAPNPC-PPPPPST    284
Chlamydomonas           ------------------------SAGGRAPVGPPLPALGEVVELEV---FE-PPDSFEV    254
Volvox                  ------------------------PGSRRAVVA-QKMRVGDEVELEV---FD-PPGSFEV    255
Ostreobium              --------------------------------------GQGDVLLEP---CE-PPADFEV    278
Chlorella               ----------------------------------QNSVEPPLVLLSC---FE-PPLDFVI    257
Picea                   -----------------------------LQLNGIDVASYQLAWLED---YQ-PPDDFVV    227
Cryptomeria             -----------------------------LQVEGINIPGHQLAWLEE---YQ-PPEDFVV    242
Taxus                   -----------------------------LQVNETNVPSHLFAWLEE---YQ-PPEDFVV    247
Triticum                -----------------------------ILWDRTR--ESEFVWLEE---YQ-PRDNFVI    226
Oryza                   -----------------------------VLPGEIN--SAEVIWLEELEDHQ-PPEGFVV    269
Zea                     -----------------------------VLPAEID--SSQVVWLEELEDHQ-PPEGFVV    234
Arabidopsis             -----------------------------LEVDGEV--VTDVTWLEE---FQ-PPDDFVI    229
Camelina                -----------------------------IEVDGEV--VTDVTWLEE---FQ-PPDDFVI    229
Nicotiana               -----------------------------LEVDGVT--FSDVTWLEE---VQ-PPDDFVV    227
Solanum                 -----------------------------LEVDGVT--FSDVTWLGE---FQ-LPDDFMI    227
Cannabis                -----------------------------LELDGVT--IPEVTWLEE---HQ-PPDNFVI    227
Glycine                 -----------------------------VQLNGVT--VSEVTWLEE---FQ-PPDNFVI    227
Medicago                -----------------------------LELDGVT--VSEVTWLEE---FQ-PPDDFVI    227
Klebsormidium           ------------------------------RDHKSSAVTDESIWLEE---FS-PPDDFVI    238
Adiantum                ------------------------------SIEDESTDGIDLAWLEE---CQ-PSDDFIV    262
Selaginella             --------------------------------EATDSAEAHPVHLQE---FR-PSSEFVV    222
Ceratopteris            --------------------------RSLTEEQRKELSLHDLVWLEE---SR-PPDDFVV    251
Ceratodon               ------------------------------------AEPGLVGWLEE---FH-PPDDFVV    241
Physcomitrium           --------------------------------VEASDQPGSIEWLEE---FH-PPDDFVV    241
Diphasiastrum           --------------------------DSTESEQELDLQEPIYVWLEE---SQPSSDDFVI    244
Marchantia              -------------------------PGGIIANGEHREESQVETMLEE---YL-PPDDFVV    242
```

```
Sphagnum                  ------------------------GSDGLQENGDNAAENPNYEWLEG---FQ-PPEDFVV      249
Caenorhabditis            -------------------------------------VAGKSAILNR---IR-CPSDFYC      224
Anopheles                 -------------------------------------DERPRYVLET---IP-NPDHYYC      238
Drosophila                -------------------------------------AEGSQVQLHC---IP-SPDTYYC      211
Candidula                 -----------------------------------KTSIGGTTWLKE---IP-QPSFYWC      229
Octopus                   -----------------------------RPLSQPEESSQKISWLVQ---IQ-QPPDFWC      239
Fasciola                  DLSSLTEAGSDVKC----ASNHQHSDSNDVQRVNGLSDGSQLIYMVE---TN-QPKDYWC      322
Schistosoma               SQNNLHNNNNNEEL----SSS---ADSSDPDLCSNQSEKDSVVYLVE---TN-QPSDYWC      304
Hydra                     LK-----------------------EYNEMLNLEEKLNLEDLTWLTE---VP-TPLDYYC      230
Daphnia                   -------------------------------------SNSEIRLLKK---TS-CPPSFWC      232
Amphimedon                -------------------------------------QDENLIWLIS---GQ-NPSWFSC      211
Lamellibrachia            ---------------------------------QGASNLEGIVCLIE---TA-QPHDFYC      235
Strongylocentrotus        -------------------------------------SESDPHWLMC---IS-QPHDFWC      243
Asterias                  -----------------------------------KETPVDLRWLLR---IP-QPSSFWC      238
Scyliorhinus              --------------------------S------PAPSPVQEAVWLLE---VP-QPSDFWC      240
Xenopus                   --------------------------GPAASDNCTGGAAQKEMWLLE---IP-QPDDFWC      238
Carassius                 ---------------------------------KSEAQEQGDLWLME---IP-QPSEFWC      243
Danio                     ---------------------------------KSEVQDQGDVWLME---IP-QPSEFWC      237
Chelonia                  ---------------------------------ADPHALPREVWLLE---TP-QAADFWC      264
Homo                      ---------------------------------SSACDLPREVWLLE---TP-QADDFWC      258
Mus                       ---------------------------------GSACDLPREVWLLE---TP-QADDFWC      244


Chytriomyces              TDGDSKSPKVPAFSKGESHKTGALKNHSSSLFEEDHEVNSEGLQMRLVELDYGDEIESVA      369
Batrachochytrium          ------------------------------------------------------------      123
Rhizoclosmatium           ----SQNSNSSTVVSTFNQHYCRKGALVSPTNSAKSFDDMCSAAKCIHESNGGSNASFCQ      327
Blyttiomyces              PRMSSSLPSLSEDDDDADEEADSDSSGSPRGTSAHPAFDPSLDALDLSACEDDADSDLVW      320
Spizellomyces             PTLDLTPPPVRSSN----------------TTISNSSSEAS-----LDLLNTSGTDEMCW      323
Chlamydomonas             VSGLYPGDPVS-------------------------------------------------      265
Volvox                    VSGCYPGDPVV-------------------------------------------------      266
Ostreobium                QRGEYRGVVIEG------------------------------------------------      290
Chlorella                 NQGRYRGVTPRCKLLPHSGSMSGVSGGDQSPTTSVHTAGGQEGHREPLLQQPSSPLSSPS      317
Picea                     QRGLYRGPKVVP------------------------------------------------      239
Cryptomeria               QRGLYKGQKVVL------------------------------------------------      254
Taxus                     QRG---------------------------------------------------------      250
Triticum                  RRDLYTGPTLEL------------------------------------------------      238
Oryza                     ARGLYKGPVIRR------------------------------------------------      281
Zea                       ARGLYKGPVIRR------------------------------------------------      246
Arabidopsis               RRIPYRGPVIRT------------------------------------------------      241
Camelina                  RRVPYRGPVIRT------------------------------------------------      241
Nicotiana                 RRGQYRGRVIKT------------------------------------------------      239
Solanum                   RRGQYKGRVIKT------------------------------------------------      239
Cannabis                  RRGLYKGPVIRT------------------------------------------------      239
Glycine                   RRGLYRGPVIRT------------------------------------------------      239
Medicago                  RRGLYRGPVIRT------------------------------------------------      239
Klebsormidium             ERGLYRGPKVQGVR----------------------------------------------      252
Adiantum                  DGAPYSGPRIIP------------------------------------------------      274
Selaginella               HRGNYRGPKVVPSI----------------------------------------------      236
Ceratopteris              TPAPYTGPTIIP------------------------------------------------      263
Ceratodon                 QRGLYRGPRIISSYN---------------------------------------------      256
Physcomitrium             QRGIYRGPRIIPSRN---------------------------------------------      256
Diphasiastrum             QEGFYKGPKIKP------------------------------------------------      256
Marchantia                QRGEYRGPRVGP------------------------------------------------      254
Sphagnum                  QRGIYRGPKVVPSMSGSNMVTTPALWNKSLRRRSTMLASHMRTRYELKLSKAVLGSEDNA      309
Caenorhabditis            DIMDFPSFEKFD------------------------------------------------      236
Anopheles                 DTVHYTPPGFMDYEEGTAPPPPPYESTTDRPV----------------------------      270
Drosophila                DVVDTPESVMQAAFQCADEVYADNASGTQ-------------------------------      240
Candidula                 DTIKYKGPLLTCL-----------------------------------------------      242
```

```
Octopus               DESDYNGPEIPSH-----------------------------------------------    252
Fasciola              ESAEYVGPSVV-------------------------------------------------    333
Schistosoma           ESAEYNGPSIV-------------------------------------------------    315
Hydra                 NTLEYTGPTFSVKS----------------------------------------------    244
Daphnia               DSLTYCGPDIKHLQHELN------------------------------------------    250
Amphimedon            IPQPYNGPKVD-------------------------------------------------    222
Lamellibrachia        DQHTYEGPEIQPVWFENDSTDSS-------------------------------------    258
Strongylocentrotus    DEVHYPGPEISIPS----------------------------------------------    257
Asterias              HQQSYLGPPVNLGQL---------------------------------------------    253
Scyliorhinus          GGEPYPGPKVVPDREAGQME----------------------------------------    260
Xenopus               GGEAYTGPKVSV------------------------------------------------    250
Carassius             GGEPYPGPEVSL------------------------------------------------    255
Danio                 GGEPYPGPKVTL------------------------------------------------    249
Chelonia              GGEPYPGPKVSP------------------------------------------------    276
Homo                  EGEPYPGPKVFP------------------------------------------------    270
Mus                   EGEPYPGPKVLP------------------------------------------------    256


Chytriomyces          YVGQQVKRDQLERGSLLTDAELVQLGGRVAMMVDRMSASTQPGQGNPSGK----------    419
Batrachochytrium      ------------------------------------------------------------    123
Rhizoclosmatium       LQVDPSGEMEVTEKEYQGLPVRDFLKLLEIAHLNGKDRVVETLKICDRIGKTFDRLVASL    387
Blyttiomyces          LAEAPDVDYDCVERAYMGPRI--FARDMIVDASDEEIYVLADALLAGQILGASGPAGGGE    378
Spizellomyces         LVEDPDVDYRCVERLYRGEPV--AKVE-----------RYANVELGII------------    358
Chlamydomonas         ------------------------------------------------------------    265
Volvox                ------------------------------------------------------------    266
Ostreobium            ------------------------------------------------------------    290
Chlorella             SAATTHSGRSGGGAQLERSNLIEAPSS---------------------------------    344
Picea                 ------------------------------------------------------------    239
Cryptomeria           ------------------------------------------------------------    254
Taxus                 ------------------------------------------------------------    250
Triticum              ------------------------------------------------------------    238
Oryza                 ------------------------------------------------------------    281
Zea                   ------------------------------------------------------------    246
Arabidopsis           ------------------------------------------------------------    241
Camelina              ------------------------------------------------------------    241
Nicotiana             ------------------------------------------------------------    239
Solanum               ------------------------------------------------------------    239
Cannabis              ------------------------------------------------------------    239
Glycine               ------------------------------------------------------------    239
Medicago              ------------------------------------------------------------    239
Klebsormidium         ------------------------------------------------------------    252
Adiantum              ------------------------------------------------------------    274
Selaginella           ------------------------------------------------------------    236
Ceratopteris          ------------------------------------------------------------    263
Ceratodon             ------------------------------------------------------------    256
Physcomitrium         ------------------------------------------------------------    256
Diphasiastrum         ------------------------------------------------------------    256
Marchantia            ------------------------------------------------------------    254
Sphagnum              AFHHAGKTNLVTTNFSVNGST---------------------------------------    330
Caenorhabditis        ------------------------------------------------------------    236
Anopheles             ------------------------------------------------------------    270
Drosophila            ------------------------------------------------------------    240
Candidula             ------------------------------------------------------------    242
Octopus               ------------------------------------------------------------    252
Fasciola              ------------------------------------------------------------    333
Schistosoma           ------------------------------------------------------------    315
Hydra                 ------------------------------------------------------------    244
Daphnia               ------------------------------------------------------------    250
```

```
Amphimedon              ------------------------------------------------------------      222
Lamellibrachia          ------------------------------------------------------------      258
Strongylocentrotus      ------------------------------------------------------------      257
Asterias                ------------------------------------------------------------      253
Scyliorhinus            ------------------------------------------------------------      260
Xenopus                 ------------------------------------------------------------      250
Carassius               ------------------------------------------------------------      255
Danio                   ------------------------------------------------------------      249
Chelonia                ------------------------------------------------------------      276
Homo                    ------------------------------------------------------------      270
Mus                     ------------------------------------------------------------      256


Chytriomyces            ----- 419
Batrachochytrium        ----- 123
Rhizoclosmatium         VVAQS 392
Blyttiomyces            HQGQR 383
Spizellomyces           ----- 358
Chlamydomonas           ----- 265
Volvox                  ----- 266
Ostreobium              ----- 290
Chlorella               ----- 344
Picea                   ----- 239
Cryptomeria             ----- 254
Taxus                   ----- 250
Triticum                ----- 238
Oryza                   ----- 281
Zea                     ----- 246
Arabidopsis             ----- 241
Camelina                ----- 241
Nicotiana               ----- 239
Solanum                 ----- 239
Cannabis                ----- 239
Glycine                 ----- 239
Medicago                ----- 239
Klebsormidium           ----- 252
Adiantum                ----- 274
Selaginella             ----- 236
Ceratopteris            ----- 263
Ceratodon               ----- 256
Physcomitrium           ----- 256
Diphasiastrum           ----- 256
Marchantia              ----- 254
Sphagnum                ----- 330
Caenorhabditis          ----- 236
Anopheles               ----- 270
Drosophila              ----- 240
Candidula               ----- 242
Octopus                 ----- 252
Fasciola                ----- 333
Schistosoma             ----- 315
Hydra                   ----- 244
Daphnia                 ----- 250
Amphimedon              ----- 222
Lamellibrachia          ----- 258
Strongylocentrotus      ----- 257
Asterias                ----- 253
Scyliorhinus            ----- 260
```

```
Xenopus                 ----- 250
Carassius               ----- 255
Danio                   ----- 249
Chelonia                ----- 276
Homo                    ----- 270
Mus                     ----- 256
```

**Supplementary Table 10 – Multiple sequence alignment of selected NCO sequences using CLUSTAL Omega.**

```
Amphimedon         MGVVKMSGLVHQLVHQARRTFSRE--------------------KTLPEEMEKLKKIMSQ       40
Homo               MPRDNMASLIQRIARQACLTFRGSGGGRGASDRDAASGPEAPMQPGFPENLSKLKSLLTQ       60
Spizellomyces      -MSPPCINESDSSPTTCCFPFCTEMENRR------------TQSLMDMSHLQRIGVLAYQ       47
Arabidopsis        -----MPYFAQRLYNTCKASFSSDG-------------------PITEDALEKVRNVLEK       36
Klebsormidium      -----MASKVQKLYDACRAAFGAKG-------------------PASKENLQLLQEALDS       36
Marchantia         -----MMTAVQRLHDVCKATFTMSA-------------------PPSAQALQRVRTILDS       36
                             .     .   *  .                        . :. :     .

Amphimedon         ITPKDFNLDSTAIDKPWDYPPFV---------------TDAPAAFMNVYECSEFNVAIFM       85
Homo               LRAEDLNIAPRKATLQ---PLPP---------------NLPPVTYMHIYETDGFSLGVFL       102
Spizellomyces      VFAAFPGRHPEADAILDRLCSLIRDVTLGHLKVKP---AQYPVEYYPIYESSVYTMCVFV       104
Arabidopsis        IKPSDVGIEQDAQLARSRSGPLNERNGSNQ--------SPPAIKYLHLHECDSFSIGIFC       88
Klebsormidium      VTCADVGLDPVTDG-ETRGFGFFGRQGPWGRHSAAPPRWAPPISYQHIYEDQHISIGIFC       95
Marchantia         IQASDVGLDEVDAMSQERGFGFFGTNGRRGRHTPMVARWAPPISYLHLYECEDFSMGIFC       96
                   :     .                                     :  ::* .  .: :*

Amphimedon         LKANKEMPLHDHPEMHGLMKILSGSMCVTSYNRPEKRGSNS-------------------       126
Homo               LKSGTSIPLHDHPGMHGMLKVLYGTVRISCMDKLDAGGGQRPRALPPEQQFEPPLQPRER       162
Spizellomyces      LKRGTTMPTHDHPKMTVFSKIVSGDLHVRTYEFAKGSTAQAHAVVDPRTG-----DLVQA       159
Arabidopsis        MPPSSMIPLHNHPGMTVLSKLVYGSMHVKSYDWLEPQLTEPE-DPSQ-------------       134
Klebsormidium      LPTCASIPLHNHPGMVVMSKLLYGDMHVRSYDWANQEAPPVGPQHPP-------------       142
Marchantia         LPTSAQIPLHNHPGMTVLSKLLYGSMHVKAYDWVNPCDEKLNSDPSV-------------       143
                   :     :* *:** *  : *:: * : :   :  .

Amphimedon         --YIARFFRS-ILATPATEPCHFTPEEGNIHQIKAQDSHVAFLDILAPPYAPEEGRDCTY       183
Homo               EAVRPGVLRSRAEYTEASGPCILTPHRDNLHQIDAVEGPAAFLDILAPPYDPDDGRDCHY       222
Spizellomyces      RPTKAGLNRIISASDPESLLLIRPEGGPNMHSFTAVSEEVVLLDIIGPPYNSE-DRPCTY       218
Arabidopsis        -ARPAKLVKDTEMTAQSPVTTLYPKSGGNIHCFKAIT-HCAILDILAPPYSSEHDRHCTY       192
Klebsormidium      -RRNAVLVCDEVLSAPCDALVLYPTSGGNIHAFTGVT-PCAVLDVVAPPYDPESGRHCTY       200
Marchantia         -PRSAKLVVNRVLTAPCETATLHPTSGGNIHAFTAVT-PCAVLDVLAPPYSAVTGRHCTY       201
                         .                     *:* : .     ..**::.***    .* * *

Amphimedon         YFKDTKQSN---------------------------------------------------       192
Homo               YRVLEPVRPK--------------------------------------------------       232
Spizellomyces      FREVSWDEGCKLVAMGVEATAPGKKSKNKKKKKRRRAVGPERKTDAMPIQENGSAPNPCP       278
Arabidopsis        FRKSRRED-L--------------------------------------------------       201
Klebsormidium      YRETPVPA-M--------------------------------------------------       209
Marchantia         YREAHSVGSN--------------------------------------------------       211
                   :

Amphimedon         --------------------------------------QDENLIWLISGQNPSWFSCIPQ       214
Homo               -------------------------EASSSA------CDLPREVWLLETPQADDFWCEGE       261
Spizellomyces      PPPPSTPTLDLTPPPVRSSNTTISNSSSEASLDLLNTSGTDEMCWLVE-DPDVDYRCVER       337
Arabidopsis        --------------------PGELEVDGEVV------T---DVTWLEEFQPPDDFVIRRI       232
Klebsormidium      --------------------VGKRDHKSSAV------T--DESIWLEEFSPPDDFVIERG       241
Marchantia         --------------------PGGIIANGEHR------EESQVETMLEEYLPPDDFVVQRG       245
                                                                * .      :

Amphimedon         PYNGPKVD------------        222
Homo               PYPGPKVFP-----------        270
Spizellomyces      LYRGEPVAKVERYANVELGII       358
Arabidopsis        PYRGPVIRT-----------        241
Klebsormidium      LYRGPKVQGVR---------        252
Marchantia         EYRGPRVGP-----------        254
                    * *  :
```
